# Automated Optimization of Bacterial Tracking Pipelines with TrackMate 8

**DOI:** 10.64898/2026.02.06.704368

**Authors:** Marie Anselmet, Laura Xénard, Marvin Albert, Rodrigo Arias-Cartin, Samia Hicham, Laura Pokorny, Élodie Paulet, Julienne Petit, Kevin J. Cutler, Benjamin Gallusser, Martin Weigert, Anne-Marie Wehenkel, Giulia Manina, Ivo Gomperts Boneca, Frédéric Barras, Daria Bonazzi, Guillaume Duménil, Jean-Yves Tinevez

## Abstract

Quantitative analysis of bacterial dynamics in time-lapse microscopy requires robust tracking pipelines, yet selecting and optimizing algorithms for specific experiments remains challenging. Indeed, Microbiologists are confronted with numerous algorithms that must be carefully chosen and parameterized to achieve optimal tracking for their experiments. We present an automated methodology to determine optimal tracking configurations for microbiological applications. It is based on TrackMate 8, a novel version of the TrackMate Fiji plugin extended with microbiology-specific tools. Our approach systematically evaluates algorithm-parameter combinations optimizing biologically relevant metrics (e.g., cell-cycle accuracy, bacteria morphology) and includes: (1) integration of deep-learning algorithms (Omnipose, YOLO, Trackastra) adequate for bacteria images in TrackMate, (2) a TrackMate-Helper extension for parameter optimization, and (3) a tracking and segmentation editor for tracking ground-truth generation. We demonstrate the effectiveness of the methodology on two use cases showing its adaptability to diverse experimental conditions. This methodology enables microbiologists with a widely applicable, automated framework to optimize tracking pipelines, facilitating quantitative analysis in bacterial imaging.

## Introduction

Light microscopy is a powerful tool for microbiology studies, giving access to bacteria morphology and dynamics over a wide range of scales [1]. Quantitative data extracted from images depicting bacteria images, alone or in conjunction with other data types, proved to be paramount to address diverse questions in the field of microbiology and infection. High-resolution images in phase contrast and fluorescence microscopy can be used to identify and localize the proteins that regulate cell elongation and division [2]. Live-cell microscopy of bacteria growth can be used to investigate a competition mechanism relevant during gut infection [3]. Time-lapse microfluidic microscopy combined with fluorescent reporter strains has proven useful to directly investigate the behaviors of single bacterial cells and their lineages over space and time [4], revealing dynamic processes and molecular clocks that could not be captured with conventional bulk-cell approaches. Preexisting phenotypic heterogeneity has been reported in both model and pathogenic microorganisms, showing how it helps individual cells adapt to fluctuating environmental conditions and promotes bacterial persistence [5], [6], [7], [8]. The knowledge gained from these studies can in turn be used to target priority pathogens such as *Mycobacterium tuberculosis* more efficiently [9]. However, analyzing single bacterial cells over space and time remains challenging, especially for species that show irregular growth and asymmetric patterns of division.

Integrating the morphology and dynamics of single bacteria in a quantitative approach involves performing tracking on a time-lapse movie. Often, a tracking tool implementation will combine (1) a detection or segmentation algorithm, able to detect bacteria, and, for segmentation algorithms, get their shape, with (2) a linking algorithm, which connect bacteria objects across the movie to build tracks and lineages, potentially through several rounds of cell division. Several tracking tools were developed specifically to address bacteria dynamics. In the last decade, end-user tools like MicrobeTracker [10] written in MATLAB, its successor Oufti [11], BactImAs [12], available as a standalone tool or as an Icy [13] plugin, the Coli-Inspector tool [14] and MicrobeJ [15] within the ImageJ [16] ecosystem were published, targeting specifically the microbiology community. This first set of tools, compared in the MicrobeJ publication, strongly rely on classical image processing approaches to segment and track bacteria. They offer different trade-offs between harnessing the wide range of bacterial shape and consistent, robust object detection. They were developed for specific applications and offer best performance within their intended contexts. But they exhibit limited generalizability when applied to broader use cases.

The advent of deep learning (DL) in biological imaging [17] offered an opportunity for better robustness. Deep-learning proved successful in segmenting cells even at high density or with challenging image quality, often outperforming classical segmentation methods [18], [19]. Additionally, they can generalize well; a pretrained neural network can be fine-tuned on new images to harness different samples and imaging modalities. These appealing features resulted in the emergence of DL-based tools to segment and track bacteria during the past years. The DeLTA tool [20], first engineered in 2020 to tackle bacterial growth in mother-machine devices [21], then re-developed in 2022 to track freely-moving bacteria [22], could achieve a very low error rate in both segmentation and tracking thanks to the use of DL. As for many tools based on DL, DeLTA is a Python toolset and requires some computing expertise. The BACMAN tool [23] is a Fiji [24] plugin that offers a user-friendly interface and bridges this requirement. It was recently augmented with a new DL-based algorithm, DistNet2D [25], that can harness very high density of bacteria with negligible errors. Omnipose [26], a DL-based algorithm extending the Cellpose algorithm [19], can segment bacteria with elongated, irregular or branched morphologies. OmniSegger [27] is the successor of the SuperSegger MATLAB tool [28] made to include the Omnipose algorithm and benefit from its capabilities. Some biological applications do not require precise cell contours, and it is sometimes sufficient to detect cells as rectangular bounding boxes to assess their motility. Detection algorithms also display lower requirements to achieve good robustness and accuracy. YOLO [29] stands out in this class of algorithms, known for its good accuracy-speed trade-off. Linking algorithms also benefited from the advent of DL, with novel neural networks that can learn the motility and division patterns of cells. DeLTA and DistNet2D mentioned above combine segmentation and linking in the same pipeline and are based on convolutional neural networks. Trackastra [30] is a transformer-based linking algorithm that generalizes well to datasets from various domains, that shines with bacteria movies. Deep learning has made bacterial analysis faster, more accurate, and accessible to labs worldwide.

This survey is just an excerpt from the vast collection of algorithms available to the microbiology community.

Thanks to them, most quantitative imaging problems become tractable for automated analysis. But their number and the diversity of their implementation make it difficult to determine the optimal combination of these tools and prototyping a solution suitable for their specific challenge.

In this article, we introduce a methodology to automatically establish the optimal tracking pipeline for a specific analysis in the field of microbiology. To make our approach user friendly and widely applicable, we implemented it in TrackMate, a Fiji plugin for prototyping tracking pipelines and analyzing tracking results. This approach is inspired by the tools we previously developed mainly around cell biology and development biology applications [31], [32]. Here, we introduce a new version of TrackMate, version 8, that ships several features and novel algorithms that will greatly facilitate tracking bacteria and analyzing bacterial lineages. In the following, we first introduce these features and detail the methodology in two use cases that cover a broad range of applications in microbiology: (A) analyzing the growth of E. coli colonies, and (B) quantifying the motility of running H. pylori bacteria.

## Results

Image analysis offers today a wide range of available segmentation, detection and tracking components that must be carefully chosen and parameterized to achieve optimal tracking results in bacteria imaging. Given the combinatorial complexity of this task, we present here an automated methodology to determine the optimal tracking configuration tailored to an experimental condition. This methodology generates tailored analysis protocols and provides an estimate of their accuracy.

### Tools and methodology

To ensure broad usability, we developed our approach within Fiji [24] using TrackMate as a core tracking platform, that we customized and expanded to integrate tools tailored for bacterial imaging. TrackMate is a user-friendly tool for automated and manual tracking of cells and organelles in 2D and 3D microscopy data. It combines state-of-the-art detection and segmentation algorithms (e.g., Cellpose [19], StarDist [18], ilastik [33], Weka [34]) with flexible tracking algorithms to reconstruct object trajectories over time in a wide range of cases, from following sub-resolution fluorescent spots to building complex cell lineages. It has a modular architecture and is easily extensible by third party developers [32]. A strength of TrackMate is its interactive interface, which allows researchers to visualize, edit, and curate tracking results. It is scriptable (via Python, MATLAB, or Fiji’s macro language), interoperable and extensible, allowing developers to integrate custom detectors, trackers, or analysis tools. The previous development line introduced support for object contours in 2D and initiated the integration of modern segmentation methods [31]. However, TrackMate v7 has included only a limited selection of algorithms, and none specifically for microbiology.

A limitation of TrackMate is that innovative algorithms for cell detection, segmentation, and tracking are predominantly developed in Python, while TrackMate is written in Java. Several solutions are being elaborated to enable the interactions between tools written in these two languages. The integration of LACSS [35] relies on a server-client model, where the LACSS Python program operates as a gRPC server listening on a TCP port. The lightweight TrackMate-LACSS Java client communicates with this server, transmitting image data and receiving segmentation results in return. Appose [36], that builds upon JDLL [37], offers Java-Python interprocess cooperation with shared memory, and is used for instance in SAMJ [38]. For this work, we elected for a simple solution based on subprocesses calling the command line interface of the Python tools, from tailored Java modules that also serve as the frontend of the tools in TrackMate. To facilitate the development of such modules, we extended TrackMate with an API that automates the commonalities of creating and calling a Python tool from Java, and that accelerates the integration of new algorithms within TrackMate. We specifically integrated Omnipose [26], YOLO [29] and Trackastra [30] for the protocols presented below. Though they are not specific to microbiology experiments, we also rewrote the Cellpose integration to include Cellpose 3 [39] and Cellpose-SAM [40], and added the spot detector algorithm Spotiflow [41] and the track visualizer inTRACKtive [42] (Figure 1).

**Figure 1:**
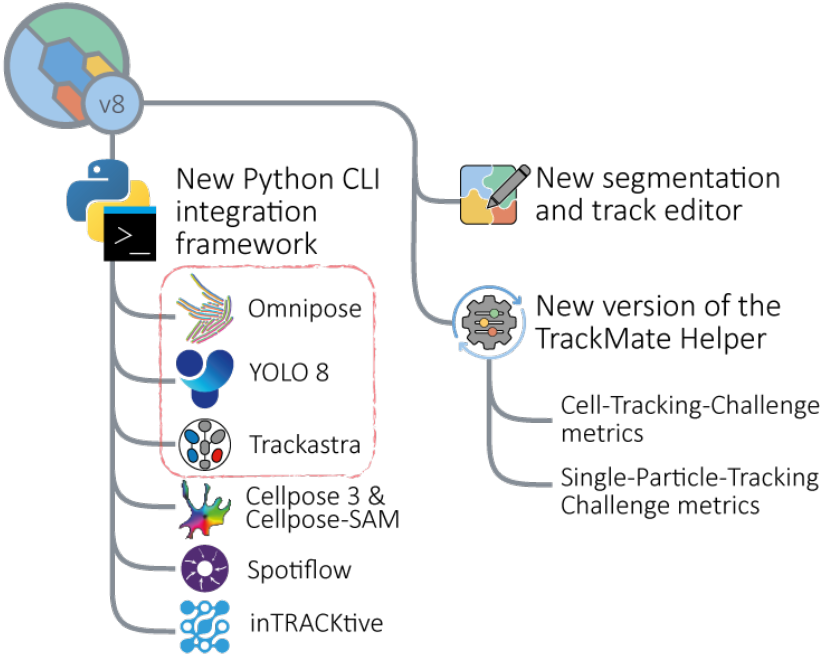
New functionalities introduced in TrackMate 8. A new framework enables TrackMate to quickly integrate external algorithms, notably in the Python ecosystem. A novel segmentation editor module based on Labkit is introduced. Coupled with the TrackScheme component of TrackMate, they allow generating tracking ground truth efficiently. A new version of the TrackMate Helper is used to perform automatic tracking parameter optimization, based on several tracking metrics that can be selected depending on the biological study. Red box: main algorithms used in the example protocols.

With these algorithms now integrated in TrackMate, we require an automated mean to optimize their selection and parametrization. To do so, we rely on tracking metrics. Tracking metrics have been initially established to rank algorithms in challenges [43], [44]. In such a challenge, a tracking algorithm runs on a movie for which the tracking ground truth is known. By comparing the algorithm output with the ground truth, several tracking metrics are computed, which are used to rate its performance, and compare it to others. Especially since the cell tracking challenge [44], tracking metrics can be linked to biologically relevant downstream analyses. For instance, the cell tracking challenge metrics include cell-cycle accuracy (CCA) and lineage branching fidelity (branch correctness, BC). They simplify assessing algorithm performance in the context of biological applications.

In our methodology, we use these metrics as objective functions for optimizing tracking parameters. First, we select an excerpt movie from a dataset and create a tracking ground truth for it. We then perform automated tracking on the movie, systematically evaluating a wide range of parameter combinations through a grid-search approach. One or two key metrics are selected based on the specific biological application and intended downstream analysis. They are used to identify the tracking configuration that maximizes them in the grid search. To automate this process and make it user friendly, we use the TrackMate-Helper extension, introduced in [31], that we fully rewrote for this work. The new TrackMate-Helper is now extensible with new metrics, includes object and track filters in the optimization and integrates the SPT metrics from [43] that are well suited for bacterial swimming. The methodology requires generating a ground truth for each dataset to analyze. TrackMate previously provided tools for manual track editing - including linking spots, removing and modifying links - but lacked the functionality to edit object contours. To address this limitation, we developed a new TrackMate module to edit object contour, based on Labkit [45]. The new segmentation editor allows to streamline track curation and object annotation in one tool and accelerates the creation of tracking ground truth (Figure 1).

In the sections that follow, we introduce this methodology for developing two distinct protocols tailored to specific use cases: bacterial growth tracking and swimming behavior analysis. We then evaluate the effectiveness of our approach and the utility of the tools introduced in this study. We first use time-lapse imaging of growing *E. coli* imaged in phase contrast to identify growth and morphology defects in mutants. In a second use case, we follow the swimming behavior of *H. pylori* imaged at high speed in phase contrast microscopy and analyze trajectories to detect how specific mutations affect the motility of the bacteria and hence, their virulence. The details of the methodology and the two analyses are given in the Supplemental Information. They follow the same process (Figure 2): (1) Generate an initial tracking results on an excerpt of the input dataset. (2) Edit this result to generate a tracking ground truth using the TrackMate manual track curation capabilities. (3) Perform tracking parameter optimization with the TrackMate-Helper, with tracking metrics adequate for the downstream analysis. (4) Perform tracking in batch on all the dataset movies with the TrackMate-Batcher, using the optimal tracking configuration found in the previous step. (5) Perform the track analysis specific to the application.

**Figure 2:**
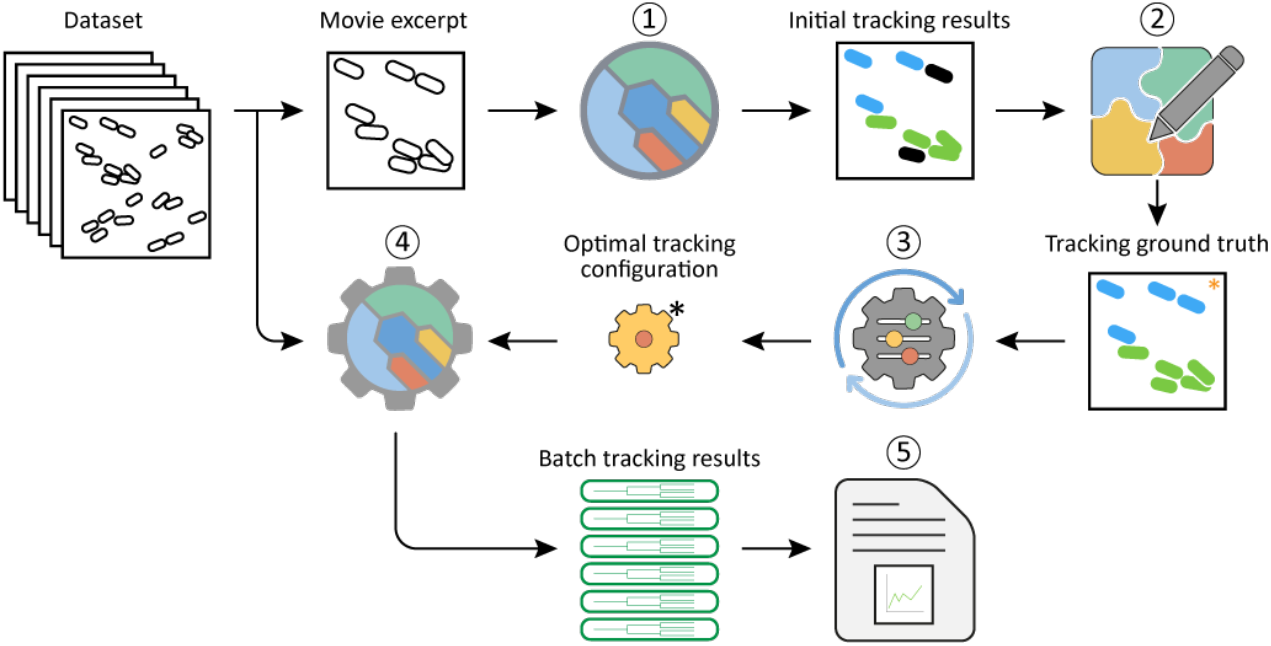
Overview of the general protocol. A movie excerpt is selected from a dataset. (1) TrackMate is used to create an initial tracking results based on sensible parameters. (2) This initial tracking results is curated with TrackScheme and the segmentation editor to generate a tracking ground truth. (3) The TrackMate-Helper is used to find a tracking configuration that optimizes selected tracking metrics, based on the ground truth. (4) This tracking configuration is then used to perform tracking in batch over all the movies of the dataset with the TrackMate-Batcher. (5) Tracking results are then analyzed with specific scripts depending on the study.

### Impact of the Ubiquinone synthesis in the growth and morphology of *Escherichia coli*

Mutations in the *Escherichia coli* respiratory chain can significantly alter cellular physiology, affecting both growth dynamics and cell morphology [46]. Quantitative analysis of these phenotypic changes provides insight into the functional roles of individual genes and can help elucidate underlying genetic pathways. Here we produce an analysis protocol that can detect and quantify growth defects in *E. coli* strains, based on phase contrast time-lapse movies. To exemplify its use, we imaged three strains of bacteria: wild-type bacteria (WT), bacteria with a mutation in the ubiquinone biosynthesis O-methyltransferase gene (UbiG), bacteria with a mutation in the 2-octaprenyl-6-methoxyphenol hydroxylase gene (UbiH). The three phase-contrast microscopy movies follow the growth of these bacteria over several cell cycles. We then performed single cell tracking with TrackMate v8 on these movies and track analysis with pycellin [47], following the protocol detailed in the Supplemental Information.

Briefly, we generated a tracking ground-truth on a movie excerpt from a 4^th^ movie following the WT strain. To speed up the ground-truth curation, we started by generating an initial tracking result from a sensible choice of tracking components and parameters. We first launched TrackMate and segmented cells with the Omnipose detector. This DL segmentation algorithm can accurately segment cells across diverse species, including those undergoing extreme morphological changes due to antibiotics or mutations. We used the publicly available model *bact_phase_omni*, which was trained on the Bacterial Phase Contrast for Instance Segmentation dataset [48]. Visual checks showed that it can segment reliably the different bacteria morphologies observed in the three movies. We then tracked bacteria with the Overlap tracker and corrected the remaining segmentation and tracking errors in TrackMate, generating a tracking ground-truth for this movie. We then performed automatic tracking parameter optimization with the ground-truth. The TrackMate-Helper tool performs a grid search for a large batch of tracking parameters and optimizes them based on specific tracking metrics. We used the segmentation accuracy (SEG) and cell cycle accuracy (CCA) metrics of the cell-tracking challenge, as we aimed at quantifying morphological and cell cycle length changes. The optimum yielded excellent scores for these two metrics (SEG = 0.904, CCA = 0.976). We then cross-validated this optimum on a movie excerpt following the *ubiH* mutant strain. We found out that, in this movie, optimal tracking configuration yielded degraded tracking results affecting the CCA value, because of spurious structures. A secondary optimization yielded a refined tracking optimum, with the addition of a filter that can remove spurious detections from tracking. With this new optimum we obtained values of SEG = 0.922 and CCA = 0.972 on the *ubiH* strain. A second cross validation with a *ubiG* movie revealed that cell cycle length cannot be measured for this mutant: most bacteria either divide once or fail to divide at all before growth arrest. In the next step we used the refined tracking optimum to batch-process all the images of the dataset in the TrackMate-Batcher. The resulting TrackMate XML files are then analyzed with pycellin.

The detailed analysis procedure is given in the Supplemental Information. Visual inspection confirms the excellent segmentation and tracking accuracy obtained with these settings, even for bacteria lineages with filamentous morphologies. The TrackMate display indicates that at late times, cell cycle length increases for the *ubiH* mutant compared to WT (Figure 3a and Supplemental Movie 1). Bacterial growth is indeed hindered for *ubiH* and even halted for most *ubiG* bacteria (Figure 3b and c). These growth defects correlate with distinct morphological abnormalities. In *ubiH*, longer cell cycles (Figure 3d) coincide with progressive filamentation, as elongated cells accumulate over time (Supplemental Movie 2). In contrast, all *ubiG* cells remain small (Figure 3e). In summary *ubiH* exhibits a prolonged cell cycle (59.8 ± 17.8 min vs. 38.6 ± 11.8 min in WT) and progressive elongation, suggesting an effect of the mutation on the division machinery. *ubiG* shows premature growth arrest after one or two divisions, with no concomitant morphological defects, hinting at an impact on a more upstream metabolic step.

**Figure 3.**
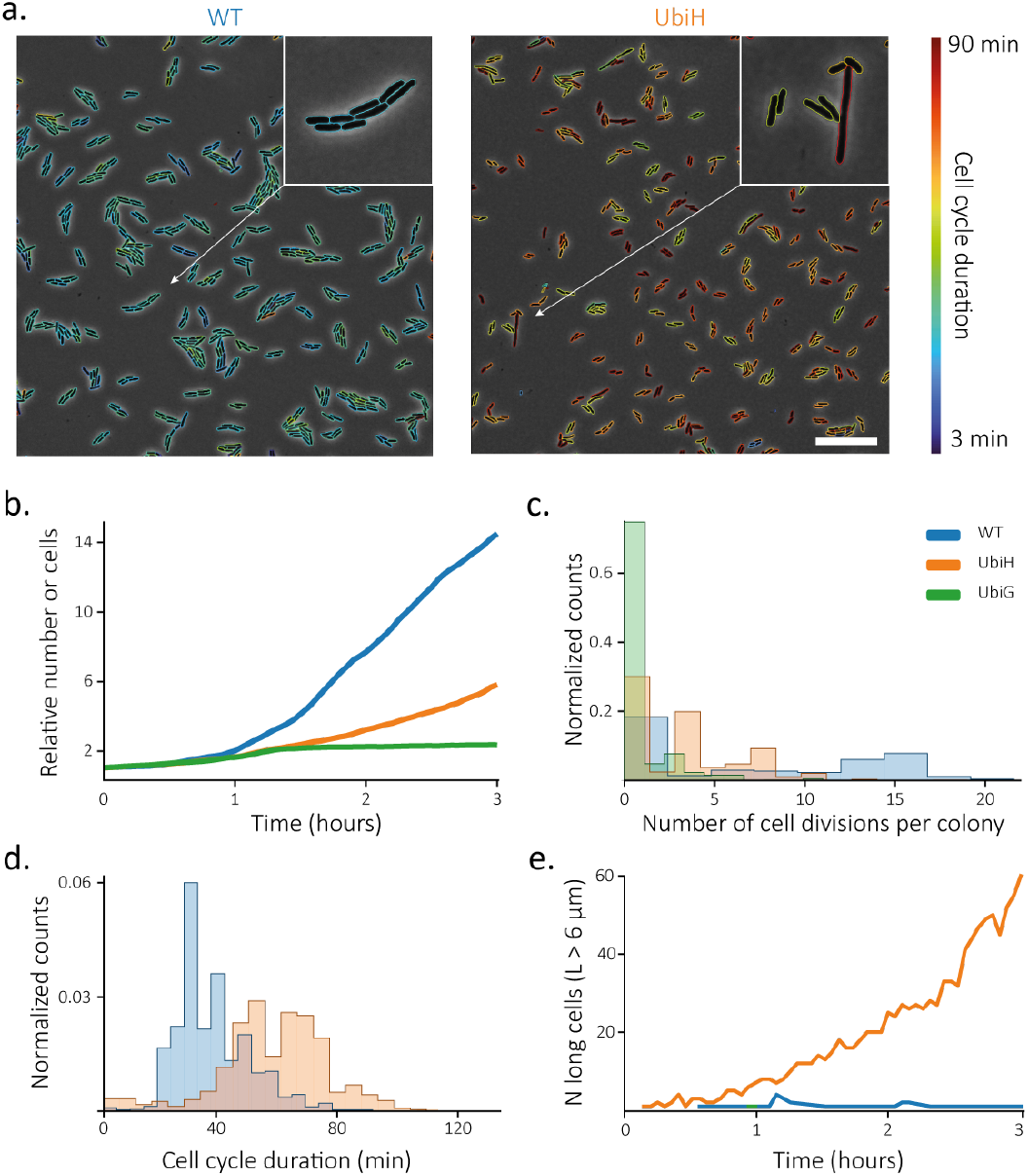
Downstream analysis of the *E. coli* tracking results.**a.**Capture of the WT and *ubiH* movies at 1 h 30 time. Bacteria are overlaid with segmentation results, colored by the length of their cell cycle. Scale bar: 20 µm. **b**. Number of cells over time in the three movies, normalized by the number of cells in the first frame. **c**. Distribution of the number of cell divisions per colony. **d**. Distribution of cell cycle length. **e**. Number of abnormally elongated cells (length larger than 6 µm) over time.

This methodology provides an automated framework for quantifying single-cell bacterial dynamics for *E. coli* imaged in phase contrast, enabling robust analysis of growth, cell cycle durations, and morphological changes. The tracking optimization scheme yields a choice of algorithms that are optimal for the specific problem at hand and gives an estimate of the expected accuracy of the downstream analysis. For instance, the tracking configuration we obtained gives us an estimate on accuracy of cell cycle length and segmentation around 97% and 90% respectively. This precision is critical for detecting biologically meaningful phenotypic differences from technical artifacts. In our case it ensures that observed variations in cell cycle length or cell elongation reflect the features of the mutations rather than tracking errors. This example also highlights the importance of cross-validation. Tracking parameters are optimized on a single movie excerpt, which must be representative of the full dataset’s morphological and dynamic variability. If not, the optimized parameters should be validated on additional movies, potentially leading to their refinements, as was the case in our study.

### Impact of *pbp1* mutations on the motility of *Helicobacter pylori*

The previous example focuses on quantifying the growth of nonmoving bacteria and their morphology. We adapt here our framework to create a protocol to quantify the dynamics of swimming bacteria and characterize the impact of mutations on bacteria motility. We use *Helicobacter pylori* as an example. *H. pylori* is a pathogenic bacterium that colonizes human gastric tissues. It is found in more than 50% of people, but only a few will develop symptoms in the shape of gastritis and ulcers. The bacteria rely on motility for its virulence [49], and an analysis pipeline that can characterize the motility of bacteria, and detect motility changes in different conditions is desirable. Such a pipeline could be used to study how genes involved in cell wall, flagella or motor proteins affect the bacteria mean speed, velocity variations, movement efficiency, etc. This requires establishing a tracking experiment where we can follow the trajectory of single bacteria, then analyze these trajectories, searching for salient differences. In the present use case, we aim to characterize how specific mutations affect the motility of *H. pylori*. We created three test groups for this use case: wild-type (WT), two punctual mutations in the *pbp1* gene involved in cell wall synthesis (mutant), and bacteria with the antibiotic selection cassette but no other mutation (control).

We seeded these bacteria in wide microscopy chambers, to avoid restricting their movement as much as possible. We then imaged them at high-frame rate and low resolution. In the resulting movies, the bacteria are moving up and down around the focus, which results in single bacteria appearing either black on a gray background, or white, or gray. In addition, their density is high, they move fast, and their trajectories are crossing each other often. Finally, the chambers contain numerous spurious dirt and aggregates that give salient signals and could confuse classical detection algorithms. All these aspects make detection and tracking challenging. To address these challenges in practice, we choose to rely on a general deep-learning detection framework, YOLO [29]. YOLO is often used on natural images but can be trained quickly to detect new classes of objects with a good accuracy. It combines object detection, classification and segmentation algorithms, but we elect to use only the detection part of YOLO. A detection algorithm only returns the position of objects it detects, not their shape. As we are only interested in the trajectory of objects this is adequate for our application. Additionally, this makes training a custom YOLO model faster and easier.

The steps of this protocol are detailed in the Supplemental Information. Briefly, we trained a YOLO model for *H. pylori* detection from a few frames extracted from the movies of the three groups. We then integrated YOLO as a TrackMate detector. In TrackMate, YOLO runs on the full movie, and the resulting detections are used for tracking. To determine the optimal tracking algorithm and parameters, we relied on the same strategy that for the previous use case. We created a small ground-truth movie from an initial tracking result, that we manually curated in TrackMate. We then used this ground-truth in the TrackMate-Helper, using this time the tracking metrics of the Single-Particle Tracking challenge [43]. Indeed, these metrics are better suited to moving objects that do not divide, and one of them, β, can be used to quantify the detection of true positive tracks and penalize spurious tracks. Unsurprisingly, the Kalman tracker performed best in this case, as it is designed for objects with nearly constant velocity. The tracking optimization also gave optimal values for the parameters of the YOLO detection and the Kalman tracker. We observed that this optimum yields some spurious detection and tracks. Several tracks are interrupted and a single bacterium can be represented by several track segments in a movie. This can be expected, given the complexity of the images. Fortunately, the subsequent analysis relies on track features. Track features are single measurements computed for each track, like mean speed, confinement ratio, etc. Their value remains accurate even with incomplete tracks. We validated that most features are not significantly different when they are measured on the ground truth tracks versus the estimated tracks. We then performed tracking with the optimal parameters on all the dataset, thanks to the Batcher, and exported the track features computed by TrackMate in CSV tables. We then analyzed the motility of the three groups based on the track features. The details of the whole analysis pipeline are given in the Supplemental Materials.

Briefly, we filtered tracks by their duration, so that the dataset includes tracks that are roughly all the same length. We first compared the WT and control groups. Differences in motility are difficult to visualize (Supplemental Movie 3) but the analysis of the tracking data can reveal whether there are track features that differentiate the two groups (Table 1). We first compared the WT and control groups. Unexpectedly, bacteria have a higher mean directional change rate (the mean speed at which a bacteria changes direction) higher in the control group than for WT (13.5 ± 5.8 rad/s vs 10.3 ± 6.0 rad/s). These values indicate that the trajectories of untransfected bacteria are straighter. This correlates with a lower confinement ratio for control against WT (0.46 ± 0.24 vs 0.60 ± 0.25), with lower values indicating that the bacteria are less efficient in going far from their starting point. Additionally, while their mean and max speed are similar, the min speed of control bacteria is lower than for WT (7.5 ± 6.9 µm/s vs 10.8 ± 10.6 µm/s). However, these differences are not significant (Mann-Whitney U test with a p-value of 0.22), and we conclude that differences in track features between the WT and control groups are not significant.

**Table 1.**
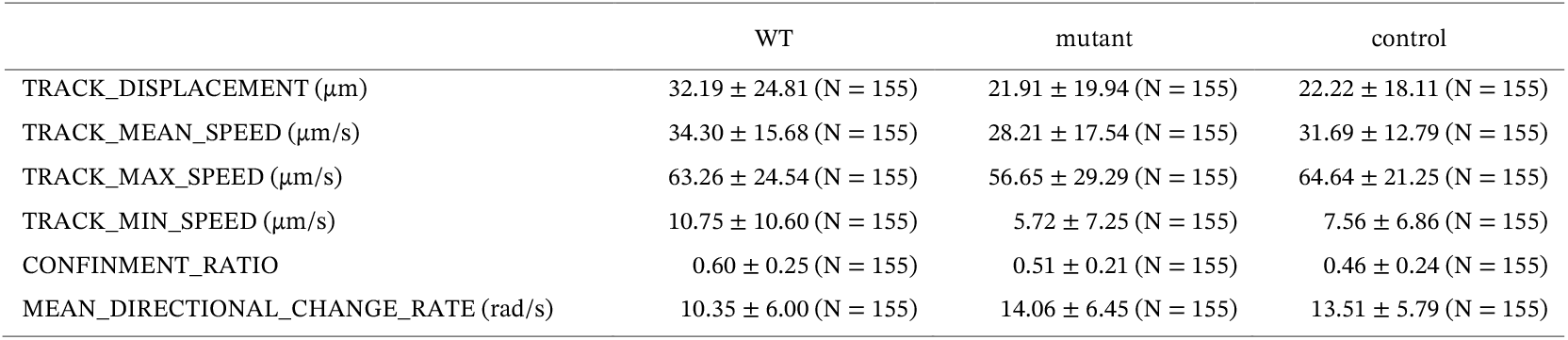
Descriptive statistics of the track features measured on H. pylori movies.

|  | WT | mutant | control |
| --- | --- | --- | --- |
| TRACK_DISPLACEMENT ($\mu\text{m}$) | $32.19 \pm 24.81$ (N = 155) | $21.91 \pm 19.94$ (N = 155) | $22.22 \pm 18.11$ (N = 155) |
| TRACK_MEAN_SPEED ($\mu\text{m/s}$) | $34.30 \pm 15.68$ (N = 155) | $28.21 \pm 17.54$ (N = 155) | $31.69 \pm 12.79$ (N = 155) |
| TRACK_MAX_SPEED ($\mu\text{m/s}$) | $63.26 \pm 24.54$ (N = 155) | $56.65 \pm 29.29$ (N = 155) | $64.64 \pm 21.25$ (N = 155) |
| TRACK_MIN_SPEED ($\mu\text{m/s}$) | $10.75 \pm 10.60$ (N = 155) | $5.72 \pm 7.25$ (N = 155) | $7.56 \pm 6.86$ (N = 155) |
| CONFINMENT_RATIO | $0.60 \pm 0.25$ (N = 155) | $0.51 \pm 0.21$ (N = 155) | $0.46 \pm 0.24$ (N = 155) |
| MEAN_DIRECTIONAL_CHANGE_RATE (rad/s) | $10.35 \pm 6.00$ (N = 155) | $14.06 \pm 6.45$ (N = 155) | $13.51 \pm 5.79$ (N = 155) |

We then assessed the effect of the punctual mutations in *pbp1* by comparing the control and the mutant groups. Here we find that the track features between these two groups are significantly different (Mann-Whitney U test 0.017). By testing for effect size, we find that differences between track feature values are all small or negligible (Cohen’s d value below 0.5 and 0.2 respectively).

The largest effect size is observed for the bacteria max and min speed, indicating that the mutations mainly affect the instantaneous speed of bacteria, albeit modestly (Figure 4c and d). From this study we therefore conclude that the two mutations affect modestly the motility of H. pylori in liquid media, by lowering the instantaneous speed by about 25% and 13% in the minimal and maximal speed (Table 1). The impact of these changes on virulence will be investigated in a follow-up study, for which this data will be valuable.

**Figure 4.**
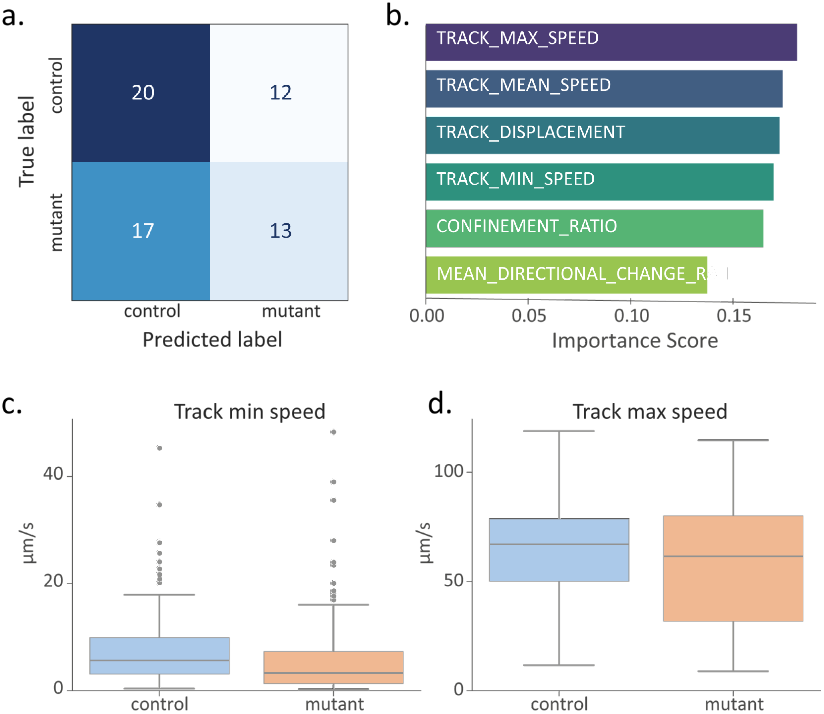
Downstream analysis of the H. pylori tracking results.**a.**Confusion matrix and **b**. feature importance of a Random Forest classifier trained against the track features measured in both the control and mutant group. **c**. and **d**. Box plot of the track minimal speed and maximal track speed respectively, measured for these two groups.

## Discussion

In bioimage analysis, a common challenge is choosing optimal algorithms in a custom-built pipeline and knowing whether it performs with a good accuracy. Our approach provides a practical solution to these questions in the context of tracking pipelines. We introduced both an updated tracking tool and a systematic methodology for determining optimal tracking parameters, relative to selected metrics and a representative ground truth. Our framework enables microbiologists to create custom protocols for analyzing bacterial dynamics across a wide range of research applications. This data-driven approach to algorithm selection greatly facilitates the analysis of time-lapse movies, which would be slowed down and open-ended in a trial-and-error approach.

The methodology relies on the integration in a common tracking platform, TrackMate, of several state-of-the-art segmentation, detection and linking algorithms. Currently, innovative image analysis algorithms are developed in the Python language, which prompted us to augment TrackMate - a Java software - with facilities to bridge to this language. The optimization itself takes place in the TrackMate-Helper, that we rewrote to be extensible and integrate new tracking metrics. These metrics were developed initially to rank tracking algorithms, but we use them to find the optimal tracking configuration for a specific problem. As expected by the diversity of imaging conditions and the bacteria that are imaged, the best algorithms differ in each case. Finally, the methodology relies also on providing a tracking ground-truth for the optimization process. This prompted another extension of TrackMate, with a convenient segmentation editor, that enables users to perform the task from end to end within Fiji.

Shipping tools as plugins in software platforms like Icy or Fiji allows them to benefit from the general features they offer. They offer first basic but key capabilities, like opening any image type, merging or separating channels, using drift correction, pre-processing, post-processing and further analysis. Secondly and often understated, these platforms offer a means to distribute and deploy plugins in a convenient manner and dealing with dependencies in a robust manner. TrackMate benefits from the Fiji ecosystem, ensuring an easy installation as well as a user-friendly interface with a lot of features available. Additionally, TrackMate ships several visualization and analysis modules that help make sense of the track analysis in the light of the image data. However, TrackMate is a generalist tool and so are its analysis capabilities. The tools mentioned in the introduction [10], [11], [12], [14], [15], [23] offer each a specific track analysis pipeline that is tailored to the application they were developed for. As we try to build protocols that remain very general, we deferred the track analysis part to bespoke scripts in Python, using the import / export and analysis facilities of the pycellin library [47]. The two use cases above demonstrate the protocol’s broad applicability across diverse imaging conditions, including variations in image quality, bacterial morphology and motility types. While the analysis is performed within TrackMate, this applicability is a tribute to the success of the algorithms TrackMate integrates. They are at the foundation of the robustness and accuracy this protocol enjoys.

The optimization procedure in this protocol can evaluate all algorithms currently integrated within TrackMate, which limits the solution space to the best-performing combination currently available there. But our primary objective is not about identifying the best algorithm: we seek tracking results of sufficient quality to enable reliable downstream analysis, regardless of the specific algorithm employed. This is exemplified in the first use case, where multiple tracking configurations yielded excellent accuracy. Nonetheless, to accommodate the growing complexity of imaging experiments in microbiology, the new integration framework of TrackMate 8 will enable rapid incorporation of new algorithms, ensuring the approach remains applicable to emerging challenges.

The tracking optimization identifies both the optimal tracking configuration and the corresponding values of the selected metrics at this optimum. These metric values provide critical insight into the suitability of the tracking results for downstream analysis, given a defined accuracy threshold. In practical cases, tracking will always be imperfect, and there will be tracking mistakes in the results even with a very good optimum. What matters is that the impact of these tracking mistakes is known, and that downstream analysis results are interpreted in the light of the tracking accuracy. In the second use case we obtained an optimum for the β tracking metrics of about 66%, indicating a substantial proportion of fragmented tracks. Consequently, we restricted our analysis to use robust track features, such as mean speed and mean directional change rate, which are less sensitive to track fragmentation. In the first use case, the high value of the cell cycle accuracy metrics (CCA value, 97%) enabled us to safely conclude that the differences in cell cycle length we observed across mutants were not the product of tracking mistakes.

In some cases, the value of tracking metrics must be interpreted with care. For instance, in the first use case the SEG metric was benchmarked against a ground truth derived from Omnipose’s initial segmentation output, which required only minimal correction. This introduces a bias in favor of Omnipose in SEG scores and makes this approach unsuitable for comparative algorithm ranking. But again, our aim is to achieve a good segmentation regardless of the chosen algorithm, and the SEG score and visual validation confirms that the optimum yields it. Importantly, this bias does not affect metrics dependent on tracking links (e.g., CCA), as all tracking errors were systematically reviewed and corrected. Thus, while SEG scores reflect high fidelity to a semi-automated ground truth, the CCA metrics provide an unbiased assessment of tracking performance. Finally, a limitation is that certain downstream analyses may require tracking metrics not included in the SPT or CTC frameworks that we used here. However, the TrackMate-Helper is extensible and will incorporate future metrics as needed.

This framework introduced here is highly generalizable. The movies in the second use case feature highly motile, non-dividing bacteria, yet the protocol-development steps remain identical to those used for the E. coli growth case. The key changes for this scenario lie in the sensible choice of an adequate object detection, in the ground truth generation, where segmentation correction was unnecessary, and the choice of tracking-optimization metrics. This demonstrates that our framework can generate robust analysis protocols for diverse experimental setups. We are also confident that it can be broadly used beyond microbiology, and possibly beyond microscopy data. This flexibility originates from two complementary aspects. On one hand, the core principles (generating a tracking ground truth, selecting meaningful tracking metrics, systematic optimization of tracking parameters) are general and not proper to microbiology or even biology. On the other hand, the framework leverages TrackMate’s highly modular and extensible architecture. It is made to be a generic tracking platform, and its extensibility enables the incorporation of new algorithms or metrics as needed. This ensures that the framework can be tailored to meet the unique demands of different fields, whether in microbiology, cell biology or beyond. Together, these two aspects enable the framework to be effectively deployed in virtually any tracking pipeline.

## Acknowledgements

We thank Chantal Ecobichon for help with H. pylori mutant construction. We thank Gaëlle Letort and the members of the Image Analysis Hub, in particular Stéphane Rigaud, for support and discussion during the project. We thank the members of F-BIAS for helping with testing the new features of TrackMate v8, and Curtis Rueden and Kevin Eliceiri for maintaining the Fiji ecosystem.

This work is supported by the Institut Pasteur, the Agence Nationale de la Recherche through France-BioImaging (ANR-24-INBS-0005 FBI BIOGEN), the European Research Council (Destop ERC AdG to GD) and the IMI 2 Joint Undertaking (JU) under Grant Agreement No 853989. The JU receives support from the European Union’s Horizon 2020 research and innovation programme and EFPIA and Global Alliance for TB Drug Development Non-Profit Organization, Bill & Melinda Gates Foundation, University of Dundee. IGB and SH acknowledge funding from the HELDIVPAT project (ANR-10-PATH-003), the Fondation pour la Recherche Médicale (EQU202403018034), the Labex-IBEID (ANR-10-LBX-62 IBEID) and the ERC starting grant (PGNfromSHAPEtoVIR, FP7-202283). LX received funding from Fondation pour la Recherche Médicale (FDT202504020138), the INCEPTION project (PIA / ANR-16-CONV-0005) and is a student from the FIRE PhD program funded by the Bettencourt Schueller foundation and the EURIP graduate program (ANR-17-EURE-0012).

## Data and software availability

The instructions and dataset to reproduce the use-cases in this article are available on Zenodo, with links and software installation instructions detailed in the Supplemental Materials.

## Declaration of generative AI and AI-assisted technologies in the writing process

During the preparation of this manuscript, the authors utilized Mistral AI to assist with language refinement. All AI-assisted content was thoroughly reviewed, edited, and approved by the authors, who assume full responsibility for the accuracy and integrity of the published work.

