## Supplemental Information for "Automated Optimization of Bacterial Tracking Pipelines with TrackMate 8"

### A. Protocol to study the impact of the respiratory chain in the growth and morphology of *Escherichia coli*

#### Overview of the procedure

We give here a detailed tutorial that reproduces the analysis described in the main text under the Results section *Impact of the respiratory chain in the growth and morphology of Escherichia coli*. In this use case our goal is to analyze a small pedagogical dataset made of three movies, covering three conditions (wild-type and two mutations), and that follow the growth of *E. coli* colonies in phase contrast. Our goal is to measure bacteria length and cell cycle length over time, and check whether we can build a protocol that can detect differences between these three groups. To determine what is a working tracking configuration for these movies, we generate a tracking ground-truth on a fourth movie that follows the wild-type strain. We then use this ground-truth in a parameter grid search, optimizing tracking metrics pertinent to the downstream analysis, namely the segmentation accuracy (SEG) and the cell cycle accuracy (CCA) of the Cell-Tracking Challenge. To validate that the optimum we find is adequate to the three movies, we check the metrics values with a tracking ground-truth built from a small excerpt of one of the three movies. We then apply automated tracking with the optimum on the three movies and then perform track analysis to characterize the impact of the mutations.

#### Install the required software

- Download and install Fiji from https://fiji.sc/. Follow the installation instructions in https://imagej.net/software/fiji/downloads

- Install three Fiji extensions. This is done in Fiji, via the *Help* > *Update* tool. Update and restart Fiji until it is up to date. Then go to the update menu once more, and click on the *Manage update sites* button, at the bottom-left of the updater window. A new window containing all the known update sites will appear. Click on the ***CellTrackingChallenge, TrackMate-Cellpose*** and ***TrackMate-Helper*** checkboxes and restart Fiji one more time.

- Install a Python distribution with conda, preferably via Miniforge:

    - https://conda-forge.org/download/

- Install Omnipose. Follow the installation instructions of https://imagej.net/plugins/trackmate/detectors/trackmate-omnipose#omnipose-installation

#### Configure conda environments in Fiji

Open Fiji, and configure the path to conda or mamba in the menu item *Edit › Options › Configure TrackMate Conda path…* as explained in https://imagej.net/plugins/trackmate/trackmate-conda-path. It is very important that the conda environments that you created are visible in Fiji.

#### 1. Generate an initial result for a tracking ground-truth

Step #1 in Figure 2.

The TrackMate-Helper can automatically find optimal tracking parameters in the sense of a given tracking metric, using a tracking ground-truth. This ground-truth takes the shape of a TrackMate movie fully tracked and manually curated. To shorten the time required to generate this ground-truth, we start from the results of automated tracking with sensible initial tracking parameters.

- Download the dataset for this use-case here: DOI 10.5281/zenodo.17909896 https://doi.org/10.5281/zenodo.17909896

- Open Fiji, and open the image in `/1-Generating an initial results for a tracking ground-truth/` named `20230331_washed_XY1.ome-1_stabilized_cropped.tif`

- Launch TrackMate from the menu *Plugins* > *Tracking* > *TrackMate*

- Click the *Next* button in the TrackMate user interface (UI). The panel '*Select a detector*' is shown. In the list, select '*Omnipose detector*' then click *Next.* If this item is not present in the list, you need to install the corresponding extension, as explained in the installation section above.

- The '*Omnipose detector*' panel is shown. We need to configure it with a few parameters:

    - The '*Conda environment*' setting is crucial. It must be the **Omnipose environment you created before**. In the image below, Omnipose was installed in an environment named `omnipose-jyt`.

    - Set the cell diameter to 3 µm. The default settings are suitable for the other parameters. You should have a configuration panel like this one:

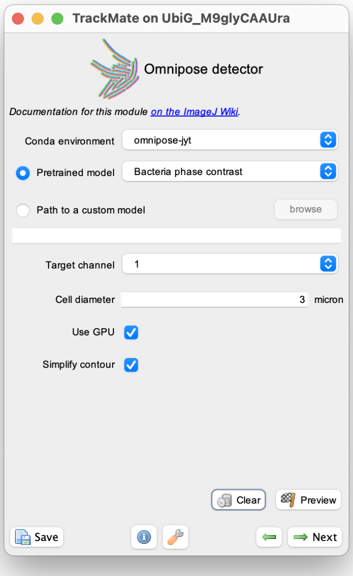

    - Verify that the configuration is working. Using the rectangle ROI in the Fiji toolbar, draw a rectangle around a small portion of the image where some bacteria are present, and click the *Preview* button. After a few seconds, the display should be updated with segmentation results, as shown below. If it is not the case, most likely the '*Conda environment'* setting is not correct, or Omnipose was not properly installed (see the previous section).

- Important: Remove the ROI by clicking outside it in the image. If there is a ROI when moving to the next step, the Omnipose segmentation will only be done within a ROI. Click *Next*. Omnipose will now run with these settings on the whole movie. Depending on your computer, this can take one to several minutes. On a Mac or on a computer without GPU acceleration, uncheck the *Use GPU* button. The segmentation will then run with multithreading on the CPU, which will accelerate computation. Once the computation is over, click *Next*.

- In the Initial thresholding panel, make sure the threshold on quality is set so that all spots are selected. Click and drag in the histogram, bringing the threshold to the left, to include all spots.

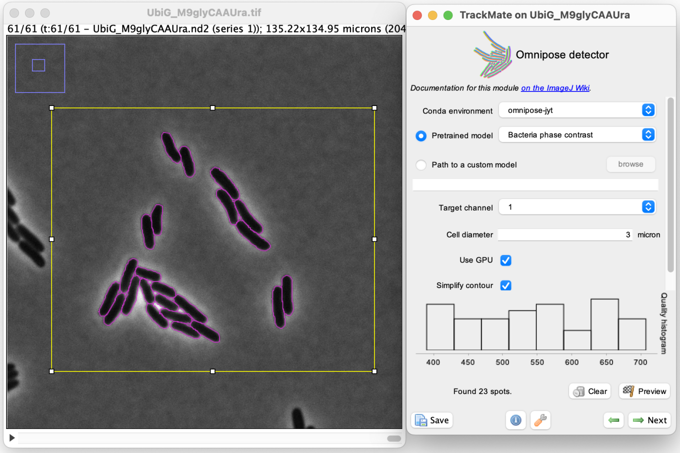

There should be almost 11k spots selected. Click *Next*.

- The *Filter spots* panel is displayed, and spot numerical features are calculated. In the image window, the segmentation results are displayed. There are some spurious detections corresponding to small fragments in the image. We can filter them out by adding a filter on their *Area*. Click on the green '+' button to create a filter panel. In the drop-down list, scroll up to *Area*. The filter box displays the histogram of the bacteria area. Click between the two peaks of the distribution to set a threshold that will filter out small fragments. A threshold value of 1.3 $\mu m^2$ is adequate. You can adjust it manually by clicking on the histogram, or by clicking anywhere in the histogram and typing the number (1.3) on the keyboard. Notice the small spots on the image disappear. The results otherwise look good and there will be few segmentation mistakes to correct. Click *Next*.

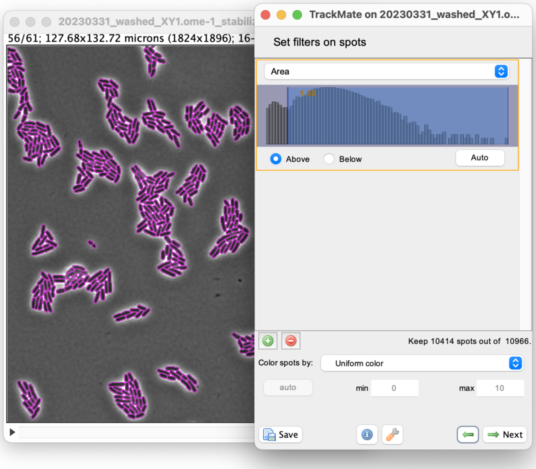

- The panel '*Select a tracker'* is shown. A reasonable choice here is the *Overlap tracker* since the bacteria do not move too much from one frame to another. Click *Next.*

- Classic settings (precise IoU calculation, Min IoU of 0.3 and Scale factor of 1.5) are good enough to generate decent initial tracking results. Click *Next.* When presented with the *Track filter* panel, make sure there are no filters there (possibly removing them with the red '-' button). There should be about 100 tracks. Click *Next* again.

- The '*Display options*' panel is the last one relevant for this step. Let's first configure the display with convenient settings. Click on the '*Edit settings*' button with a wrench in the top-right of TrackMate UI. A new panel titled *TrackMate display settings* is shown. For *spot color* and *track color*, choose '*Track index*'. Click on the *draw spot filled* checkbox and set the *spot alpha transparency* to 0.5. In the *track display mode*, select *Show tracks backward in time.* Finally scroll down to the *line thickness* setting and set it to 2. Close the panel.

- You can now save the tracking results. Click on the *Save* button at the bottom-left of TrackMate UI and point to the location next to the image file.

The TrackMate XML file that results from our run of this step has been saved in the Zenodo folder for this use case, in the folder `/1-Generating an initial results for a tracking ground-truth/results/`. This is the one we use below. To follow the next section with it, move the XML file in the same folder as that of the image, and open it in TrackMate with the command *Plugins* > *Tracking* > *Load a TrackMate file*.

### 2. Curating the automated tracking results

Step #2 in Figure 2.

There are few segmentation and tracking mistakes that need to be corrected before we can use these results in parameter optimization. For this we need to inspect results and detect mistakes first. TrackMate offers several facilities for this. The main one is TrackScheme, the lineage browser of TrackMate. Open it by clicking on the *TrackScheme* button on the TrackMate UI. TrackScheme is documented in detail here: https://imagej.net/plugins/trackmate/views/trackscheme. We describe below how to use it for detecting errors and the manual curation of tracking results in our case. Fixing tracking mistakes can be done fully in TrackMate, and requires being familiar with its interface to correct, add and remove links. This is documented here:

- https://imagej.net/plugins/trackmate/tutorials/manual-track-editing for editing links in TrackMate,
- and here https://imagej.net/plugins/trackmate/tutorials/manual-tracking for editing links inside the image.

Finally, we will need to edit faulty segmentation results, which can be done with the segmentation editor of TrackMate, based on Labkit. It is also documented online here: https://imagej.net/plugins/trackmate/tutorials/trackmate-segmentation-editor. The paragraphs below repeat nonetheless some of the information found in these four tutorials in the scope of this use case.

Most of the time, linking and segmentation errors manifest in obvious tracking mistakes that can easily be detected with TrackScheme. Indeed, for this kind of bacterial dynamics where bacteria divide rather regularly without moving a lot, we expect the lineages to resemble binary trees, where each branch divides into two daughter branches until the end of the movie. When a lineage deviates from this pattern, this indicates potential tracking mistakes. We give here a few examples for most common situations and how to fix them.

#### Fixing over-segmentation mistakes

In the movie, at position X = 63 µm and Y = 50 µm a bacterium is over-segmented twice, at frames 34 and 36. This is caused by the bacterium displaying a small dip at its equator, which the Omnipose segmentation model mistook for a complete division. This results in having a lineage tree with dangling branches, as shown below. Fixing this error will involve merging the two fragments of the over-segmentation events in one. We detail below the steps needed to fix these two errors specifically.

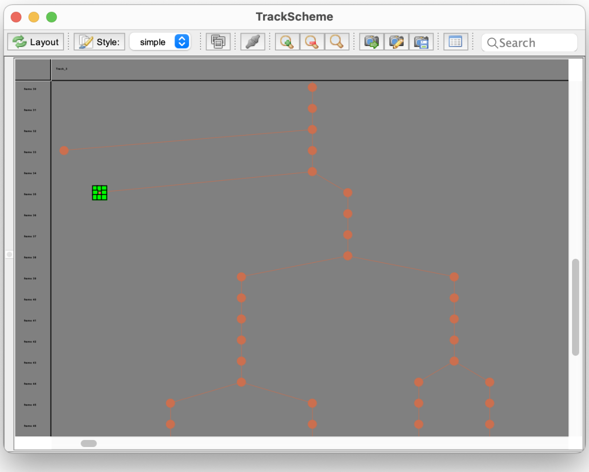

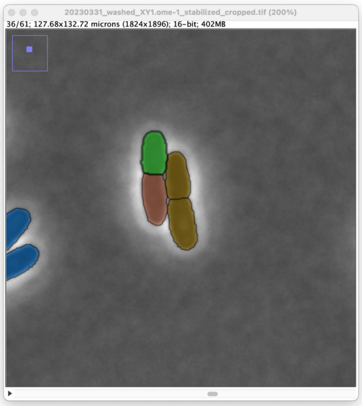

Let's start with the one at frame 34. Set the image display to this frame and zoom in around the location to fix. We can hide the track display by unchecking the *Display tracks* button in the TrackMate panel. To simplify editing, we can open the spot editor on just the region of interest. To do this, draw a rectangular ROI around the objects to merge. The spot editor will use the same color scheme for objects than the one used in the image display. Since by default it colors objects by track ID, it makes it difficult to visually differentiate the two fragments. In the *Color spots by* list, select *Random color*.

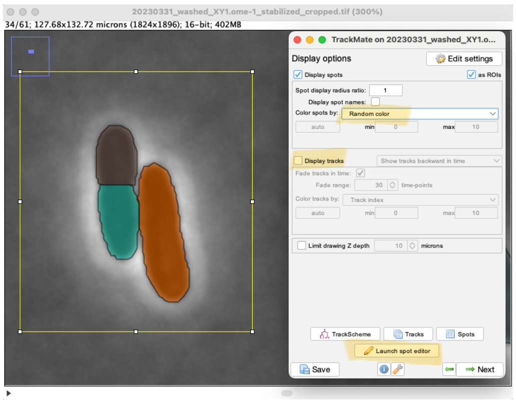

Then press the *Launch spot editor* button. The editor opens with only the data from the currently displayed frame, within the ROI if any.

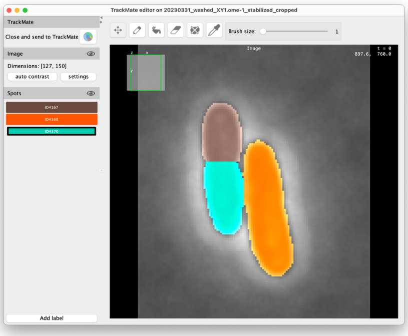

This is a Labkit window, stripped down to display only the label editing tools. We want to merge the brown fragment with the cyan one, which is done by replacing it by the cyan label in the top segment. To do this quickly, select the cyan label in the *Spots* side panel (or *shift-click* into the cyan fragment). Then select the *Flood fill* tool in the top toolbar (the one that resembles a spilling coffee mug), make sure that the *Flood mode* is set to *Replace* and click inside the brown label.

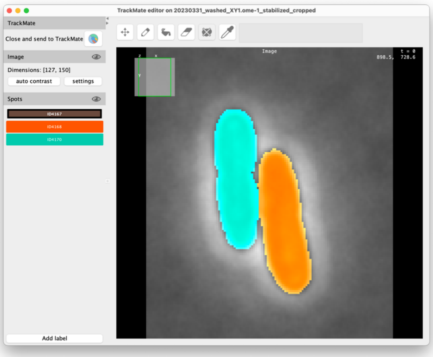

Then close the Labkit window. A dialog asks you whether you want to commit these changes to the TrackMate results. Click *Yes* and leave the *Simplify contours* box checked. TrackMate now updates the lineage with the corrected object. Note that for TrackMate, this correction involves removing the two small fragments and adding a new larger object instead. TrackMate tried (here successfully) to reintroduce the new object in the lineage with a hierarchy that is sensible given the existing links before correction. When it fails, you will have to recreate the correct links. We get the following lineage (after coloring the spots again by *Track index*):

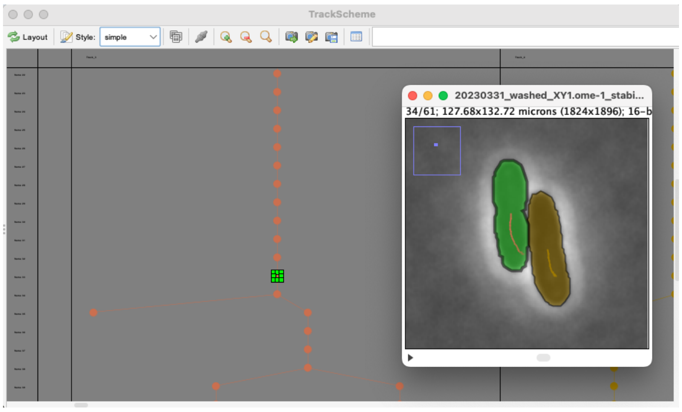

Let's now fix the over-segmentation in frame 36. Move to this frame, choose again *Random color* for coloring spots and with the ROI still present, click on the spot editor button. This time we will merge onto the isolated fragment to confuse the lineage and see what happens then.

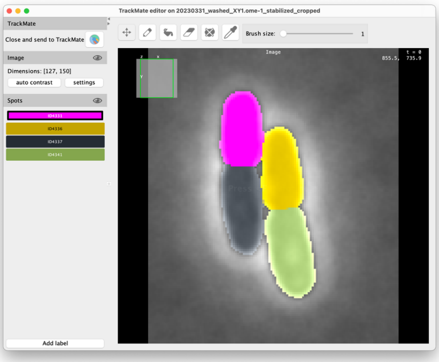

As previously, select the magenta label, pick the *Flood fill* tool (again with *Floor mode Replace*) and click inside the gray label to add the magenta one. Close the editor and accept changes. Now that we erased the object that was linked within parent lineage, the correction results in having two lineages with a break where we fixed the segmentation.

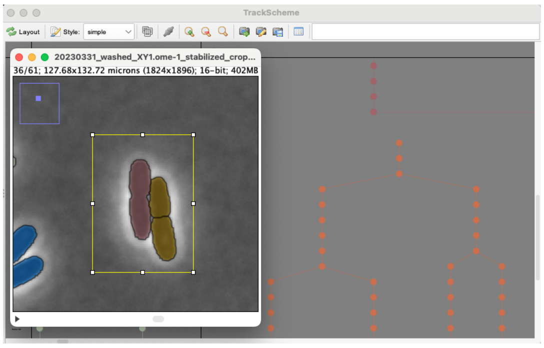

We need to link them again. To do so, make sure the TrackMate tool is activated in the Fiji toolbar (rightmost one with a magenta lineage button), not the rectangular ROI tool. Click in the image on the fixed object at frame 36, move to the next frame (shortcut: *G*) and *shift-click* into the child object to add it to the selection. Then press the *L* key to create a link between them. In TrackScheme, press the *Layout* button and the *Style* button in the top-left of the UI to rearrange the lineages. The lineage is now mended.

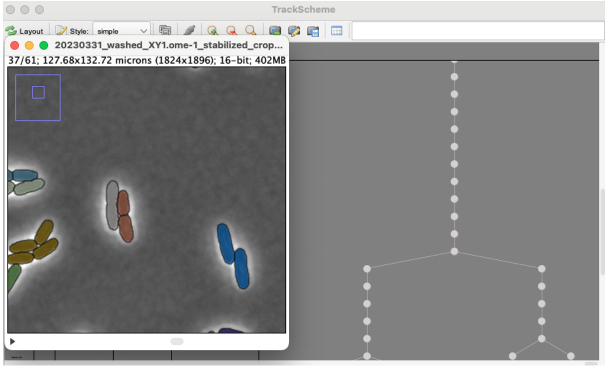

#### Fixing under-segmentation mistakes

Under-segmentation happens when a single segmentation spot overlaps with several bacteria. A salient example can be found at frame 60, at the position X = 54 µm, Y = 54 µm. In the lineages, such an event corresponds to branches that stop early. In the frame before the under-segmentation events, several objects should link to the same number of objects. But because of the under-segmentation, some are missing, which results in lineage branches to stop. Fixing this mistake will require splitting the under-segmented object into several ones, matching the bacteria. (The colors on the images below might not match what you have.)

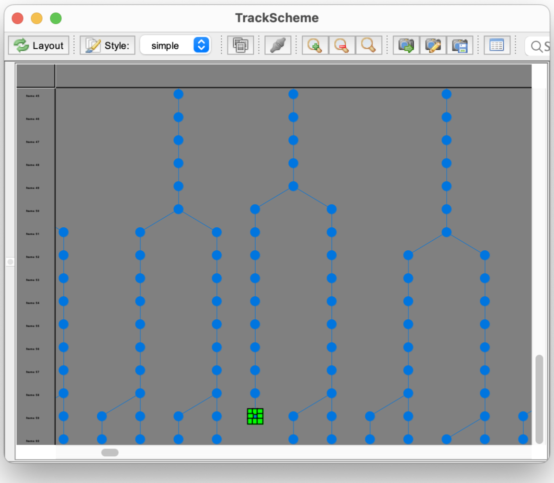

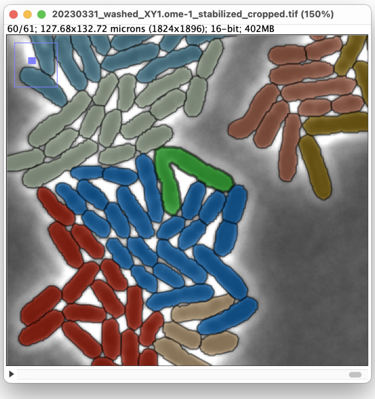

As above, this is done with the spot editor. Select *Random color* for the spot coloring, draw a rectangular ROI around the object of interest and click on the *Launch spot editor* button. To fix this mistake we will need to cut the label in half. To do this, in the Labkit editor, first select the label of the spot to edit (the purple one below, ID12200), then select the *Erase* button (the one that resembles a pen eraser) and draw a line to separate the wrong label in two. If the *Erase mode* is set to *All labels*, the pixels will be erased no matter the selected label. With the *Selected label* mode, you need to select the label that you want to edit in the left sidebar.  You may want to switch the visibility of either the image (the eye button next to the *Image* sidebar) or the labels (next to the *Spots* sidebar) to ensure the label reflects the actual contour of the bacteria.

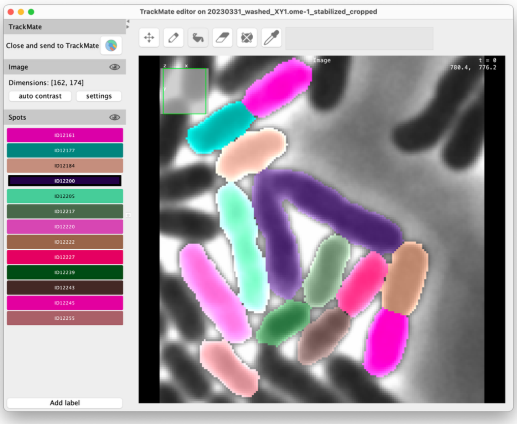

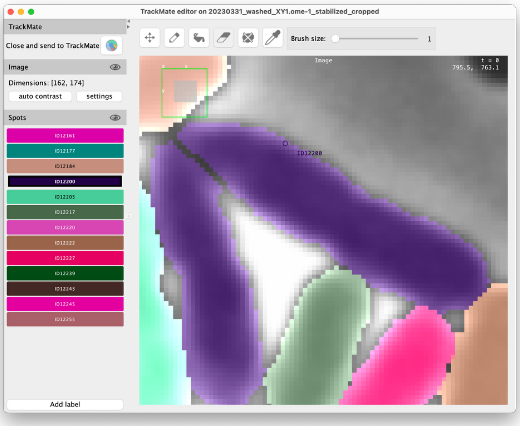

The two now separated fragments still have the same label in the spot editor. However, as soon as they are separated by at least one pixel everywhere, TrackMate will interpret them as two separate spots. If you need to create two separate objects that are touching, first separate them as explained above, create a new label (*Add label* button at the bottom left) and paint one of the two fragments. Then you can add pixels with the new label selected with the *Draw* tool (that looks like a pen) until the two fragments are in contact. If they have different labels (even with the same color), they will generate two spots in TrackMate.

After closing the editor and accepting the changes, the image display resembles this:

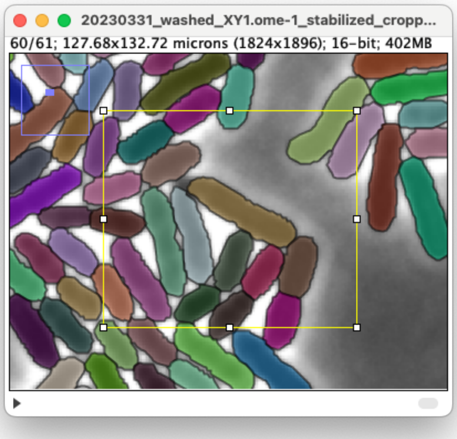

You must now relink one of the new fragments to the lineage. For this, proceed as explained in the previous section.

#### Fixing false-negative link

A false-negative linking event happens when a link between two objects in two consecutive frames was not found by the linker. This happens for instance at frame 58, at position X = 19 µm, Y = 83 µm. The bacterium moved enough so that the overlap tracker we used missed the link. This results in having a small tracklet for the child lineage, as shown below.

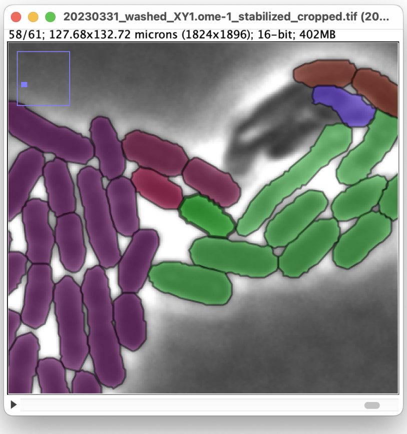

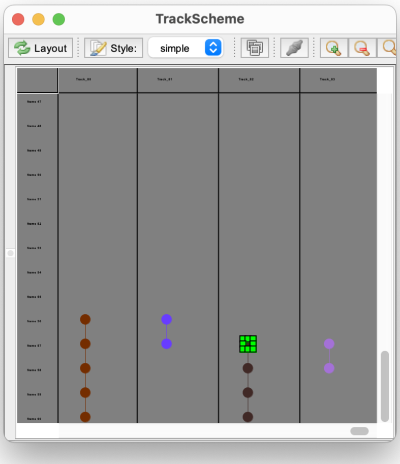

Correcting this kind of error does not require fixing the segmentation. It can be done directly in the image overlay. Move the frame 57 and click on the parent object to select it. Move to frame 58 and shift-click on the child object to add it to selection. Then press the *L* key to link them. A link is created between the two objects. In TrackScheme, press the *Layout* button in the top-left of the UI to rearrange the lineage.

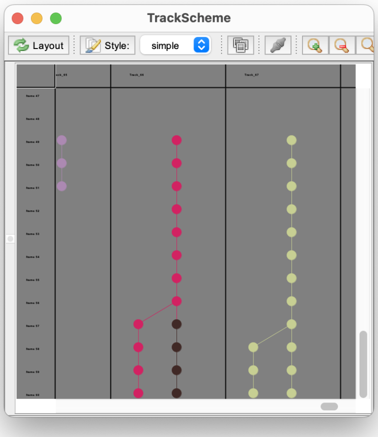

Then press the *Style* button to match the object colors between the image overlay and TrackScheme.

#### Fixing false-positive link

A false-positive link corresponds to the opposite event: a spurious link was created by the linking algorithm. This spurious link connects two objects that should be connected with other objects, so we need to modify at least two links to fix this error. Our initial tracking results have few of this type of errors. They manifest in TrackScheme by having lineages that have too frequent branching events. One of them happens at frame 53, at location X = 54 µm, Y = 49 µm.

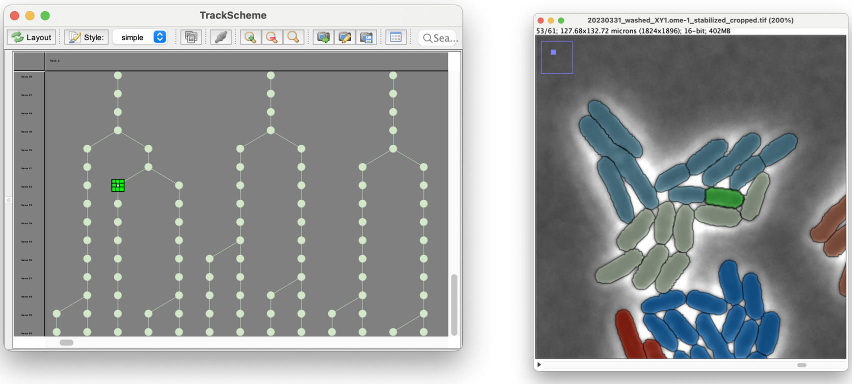

The object highlighted in green should have been linked to the lineage in blue (in the image above) but was instead linked to the lineage in ivory color. To fix it, we first remove the faulty link. Select it in TrackScheme and make sure you have the right one by checking in the image view.

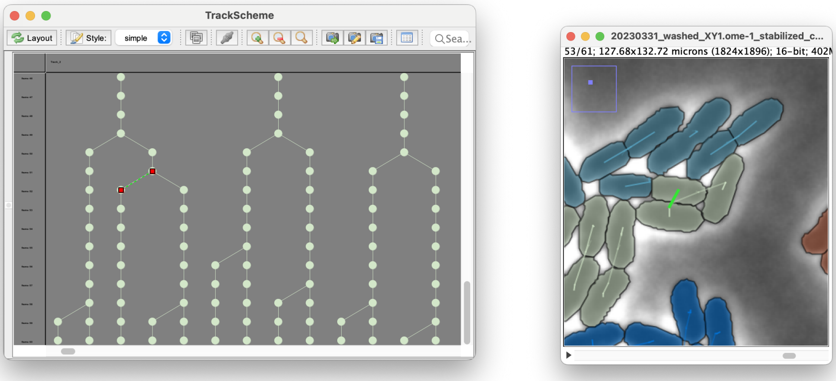

Then press the *delete* key.

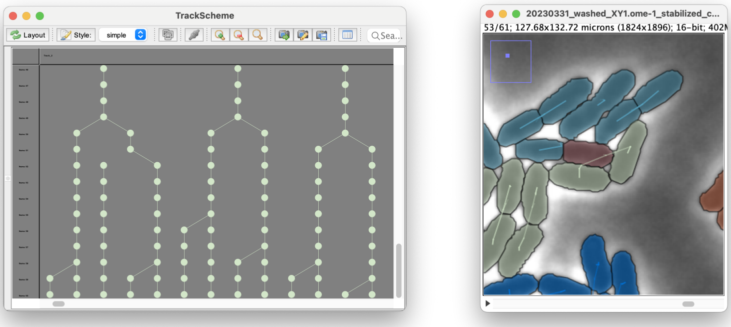

Now we need to reconnect the dangling branch to the parent lineage, via the right parent object. Click the image window to select it, and in the image, click in the bacteria object that we just disconnected in frame 53. Press the *F* key to move to the previous frame and *shift-click* into the parent bacteria object. Then press *L* to create a link between them. The blue lineage is now mended.

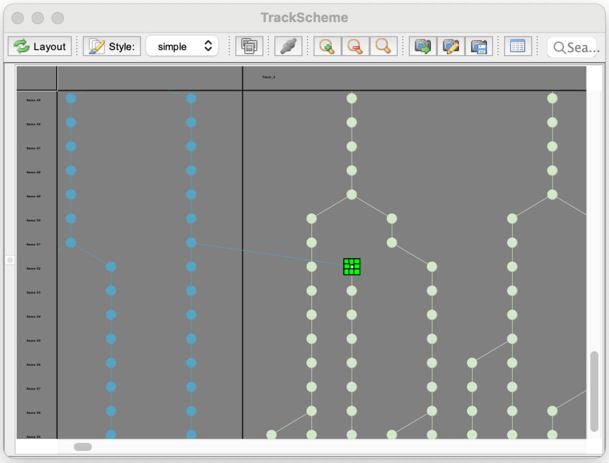

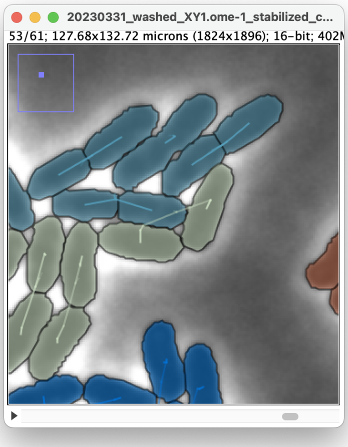

In TrackScheme, press the *Layout* button and the *Style* button in the top-left of the TrackScheme UI to rearrange the lineage.

#### Finishing curation

These four types of error are the main ones you can find when curating tracking results. They can come from segmentation mistakes (over or under-segmentation) or from linking mistakes (missing true links or creating spurious links). Importantly, they are easy to detect in the lineage view of TrackScheme. Each error will result mainly in having lineage branches that stop too early, or small branches that do not start with the beginning of the movie. You can therefore detect all errors in TrackScheme by inspecting all the lineages one by one, inspecting branches that terminate before the end of the movie, then inspecting dangling tracklets. Fixing them then involves first fixing the segmentation errors with the Labkit editor in TrackMake, then the spurious and missing links as we have explained above.

All the tracking mistakes need to be fixed before moving on to the next step. This can be long but is doable iteratively in several curation sessions (see Supplemental Table 1). To mark what lineages have been inspected and fixed already, you can rename either the whole lineage by *double-clicking* on the top row header, or by selecting several spots or all the spots of a lineage and editing their name.

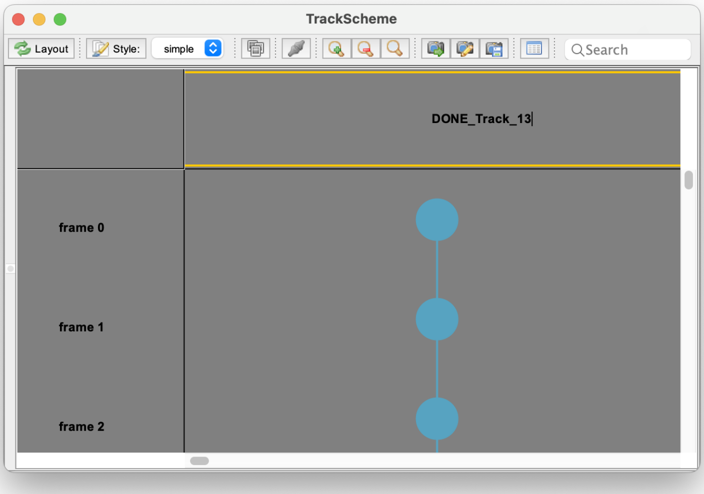

Following this method we generated a tracking ground-truth for this movie, that we have saved in the results folder `/2-Curating the automated tracking results/` under the name `20230331_washed_XY1.ome-1_stabilized_cropped-GT-DONE.xml`. Three of the 72 lineages display problematic tracking errors that are difficult to fix. They correspond to events where cells move out of focus, overlap or die, which is difficult to assess with transmitted-light imaging. We fix these errors to the best of our ability and use the results as ground-truth in the next section.

### 3. Optimizing tracking parameters

Step #3 in Figure 2.

There are mainly two factors that make it impossible to obtain perfect tracking results. First there are some segmentation mistakes that may be unrecoverable with an automated algorithm. As we noted in the section above, because of the imaging modality imposed in this study (2D, transmitted light), there are some rare situations where it is impossible to ascertain a segmentation result. Second, timelapse experiments have a limited frame rate. Since we do not have infinite imaging speed, there is a limit to the number of events we can capture. Nonetheless, we can still try and optimize our tracking pipelines to get the best results we can have with the tools at our disposal, and reason whether their accuracy will be sufficient for the downstream analysis.

#### The TrackMate-Helper

The TrackMate-Helper is a TrackMate extension that performs tracking parameter optimization. Its principle is simple: given a ground-truth tracking results, it performs tracking by iterating over a wide range of tracking parameters, and for each, measuring the accuracy of the results with respect to the specified ground-truth. It then presents the best parameter set with respect to a specific tracking metric. For this study we redeveloped the TrackMate-Helper so that it can perform optimization over a wider range of algorithms, include filters in tracking parameters, and use different tracking metrics. It can use the Cell-Tracking Challenge [1] metrics (CTC) and the Single-Particle Tracking challenge [2] metrics (SPT). The SPT metrics are adequate for objects that move in a crowded environment, but do not divide. In Life-Science they are used for organelles and quickly moving cells that do not divide. They do not include a metric on segmentation accuracy. The CTC metrics focus on cell dynamics, possibly with divisions, and include a segmentation metric. In this study we need to measure the cell cycle length, and assess the bacteria length, hence the CTC metrics are adequate.

In the folder `/3-Optimize tracking parameters with the TrackMate Helper/` you will find a copy of the TrackMate file that stores the ground-truth: `20230331_washed_XY1.ome-1_stabilized_cropped-GT-DONE.xml`. Copy the image file `20230331_washed_XY1.ome-1_stabilized_cropped.tif` there. We first need to export it to the CTC file format. Open the file in TrackMate with *Plugins* > *Tracking* > *Load a TrackMate file*. Once it is open, use the *Next* arrow in the TrackMate UI to move to the last panel called *'Select an action'*. Here there is a list of miscellaneous actions, and one of them is called *Export to CTC format*. Execute it. A dialog is shown that prompts you were to save the data. Put it in `/3-Optimize tracking parameters with the TrackMate Helper`, which will be our working folder. In the '*Data is'* list, select '*Silver truth'*. Following the CTC conventions, this indicates that this ground-truth was generated from an automated algorithm then manually curated[1]. We use this tag to stress that we started from the Omnipose algorithm, and that some metrics, notably the segmentation ones, will be biased towards it. The export will generate two subfolders, required to evaluate CTC metrics: `01` that contains the movie frames and `01_ST` that contains the tracking ground-truth.

Close the TrackMate window. Now with the image still open, run the command *Plugins* > *Tracking* > *TrackMate Helper*. The Helper launcher dialog appears. Select the CTC metrics, in the image list, make sure the bacteria image is selected, and in the ground truth path, browse to the folder generated by the export above, and named `01_ST`. Click *OK* and the Helper window opens.

The Helper is documented here: https://imagej.net/plugins/trackmate/extensions/trackmate-helper and we detail below the procedure for our use case. The window is divided into three parts. The top-left one is used to run and stop the optimization; the top-right one is used to select what components are part of the optimization; and the bottom tabs are used to check results and configure individual components.

[1] The CTC convention is slightly different however: *silver truth* indicates that the ground-truth was generated from the best automated tracking algorithms according to CTC metrics, without manual curation. In our case we manually curated the results.

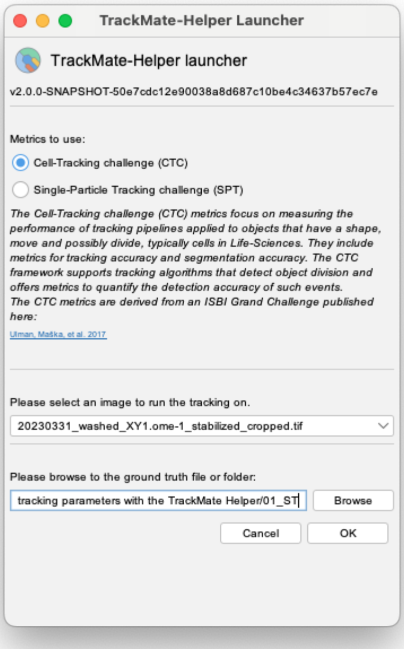

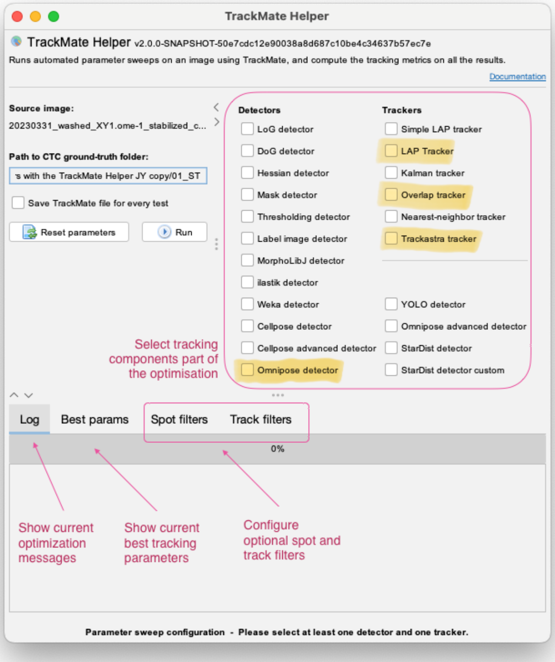

#### Configuring a parameter sweep

Let's start by selecting the components for optimization. On the *Detectors* and *Trackers* panel select the *Omnipose detector*, the *LAP tracker*, the *Overlap tracker* and *Trackastra*. A new tab appears for each of these four modules in the bottom panel. You can configure the parameters to test in the tab of a specific detector or module.

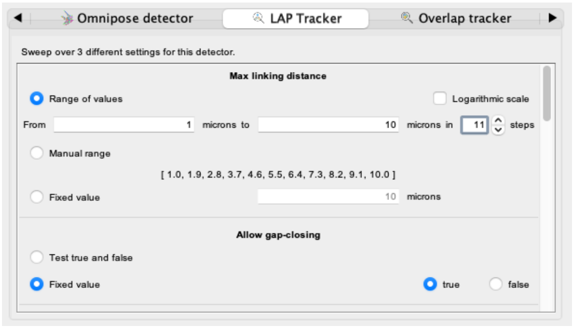

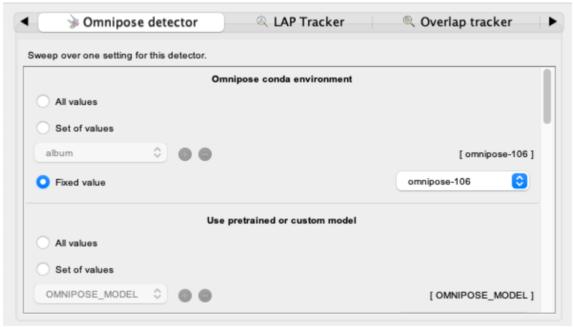

For instance, the LAP Tracker is configured with a list of parameters (*Max linking distance*, *Allow gap-closing*, etc) that are listed from top to bottom. For each of them, several widgets let you configure the range of values to test. The *Max linking distance* is a numerical value, and you can choose to test a range of values evenly spaced, on a logarithmic scale, or entered manually. You can also choose a fixed value. When the parameter expects a string of characters, like the path to the Python environment for the Omnipose detector, you can enter several of them by clicking on the green '+' button.

Selecting a parameter range is a matter of balancing between a sufficiently large parameter space and execution time. When algorithms are fast, you can test over many parameters safely. We recommend first searching interactively for sensible parameter values by using the classical TrackMate plugin running on a subset of the movie, then configuring the helper to scan around these values.

**Omnipose detector**

See this page for installations instructions: https://imagej.net/plugins/trackmate/detectors/trackmate-omnipose

For the sake of time, we will configure this detector with only one setting, the one that we have used above when preparing the ground-truth. This means that in this run, we will not optimize over segmentation parameters, but only on linking parameters.

- Conda environment: Select the conda environment in which you installed Omnipose, as when configuring TrackMate in the previous section.
- Omnipose model: Select *Fixed value* and choose *Bacteria phase contrast*
- Cell diameter: Select *Fixed value* and enter 3 µm
- Channel to segment and Optional second channel: For both select *Fixed value* and enter 0, which should be the default.
- Use GPU: Select *Fixed value.* If you have a Mac, select *false*, *true* otherwise.
- Simplify contours: Select *Fixed value* and *true*.

**Spot filters**

A quick inspection of the movies revealed that there are small dust particles that might generate false positives detection with the Omnipose detector. We can check the importance of filtering them out by adding a parameter sweep over a spot filter based on object size. In the *Spot filters* tab click on the green + button to add one filter. A panel appears to configure it

- Feature 1: use *Fixed value*, and *AREA*.
- Threshold: use *Range of values*, from 0 to 5 $\mu m^2$ in 6 steps
- Above?: *Fixed value*, *true.*

This will configure a spot filter on the object area that will test 6 different values below which to remove spots. The value 0 is used to skip filtering.

**LAP detector**

We will include the trackers that can deal with cell divisions. The first one is the LAP tracker, adapted from [3].

- Max linking distance: Select *range of values*, from 1 µm to 10 µm in 4 steps.
- Allow gap-closing: *Fixed value*, true.
- Gap-closing distance: *Range* from 1 to 10 µm in 4 steps.
- Max frame gap: *Range* from 2 to 5 frames in 4 steps.
- Allow track splitting: *Fixed value*, true. This is the parameter that lets the algorithm find cell division.
- Splitting max distance: from 1 to 10 µm in 4 steps.
- Allow track merging: *Fixed value*, false. Fusion of cells is irrelevant in our case.
- Merging max distance: Ignore.

This alone will generate 256 parameter combinations to test. Fortunately, this tracker implementation is very fast.

**Overlap tracker**

As the bacteria do not move too much in our case, we expect this algorithm to perform well with default values. Nonetheless we will explore a reasonable range of parameters.

- Scale factor: *Range of values*, from 1 to 2 in 6 steps.
- Minimal IoU: *Range of values*, from 0 to 0.5 in 6 steps.
- IoU calculation: *Fixed value*, precise.

**Trackastra tracker**

See this page for instructions on installing Trackastra: https://imagej.net/plugins/trackmate/trackers/trackmate-trackastra

The advantage offered by deep-learning tracking algorithms is that they require little parameters. In our case we will test all the models' combinations.

- Trackastra conda environment: *Fixed value*, put there the name of the conda environment in which you installed Trackastra.

- Pretrained model: *All values*.
- Use pretrained or custom: *Fixed value,* PRETRAINED_MODEL.
- Trackastra custom model path: Ignore.
- Tracking mode: *All values*.
- Use GPU: *Fixed value*, automatic.

We will not be using spot and track filters. With this configuration, these four modules should generate 1788 different tracking parameters to test. This number is simply calculated from the possible detector and tracker combination:

$$n_{total} = (n_{omnipose}) \times (n_{spot\_filters}) \times (n_{LAP} + n_{overlap} + n_{trackastra})$$

#### Running the parameter sweep

Select the *Log* tab and click the *Run* button in the top right panel. Some messages will appear in the log, reporting the progress of the parameter sweep. The helper will start by running the Omnipose detector on the whole movie, which will take time. Then it will iterate over all the tracking parameters, without re-running the detection step. Once the first tracking run has completed, the results will be evaluated (using the CTC metrics), and the *Best params* tab should display a table reporting the current optimization results. It will be updated live as the optimization progresses. The full run might take a few hours depending on your workstation. It can be run as an overnighter. If you must stop the optimization, you can relaunch the helper by pointing it to the same ground-truth folder and input image. It will remember the parameter sweep you configured and will resume the optimization from the last completed run.

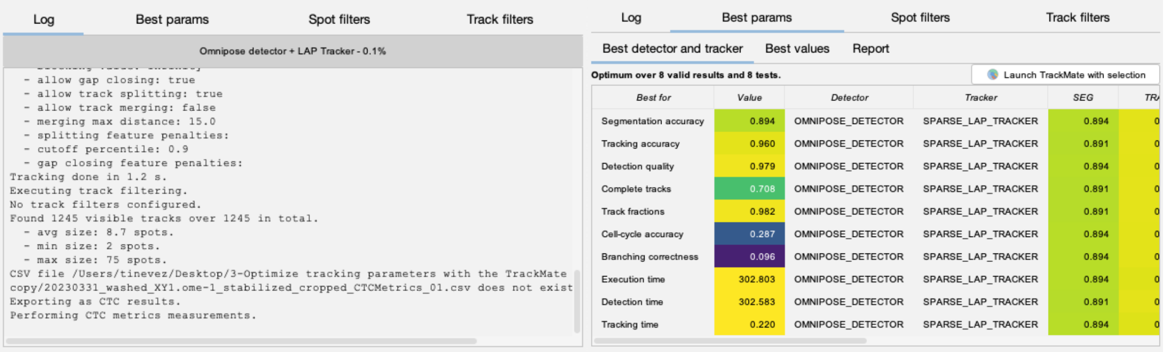

#### Interpreting optimization results

The folder `/3bis-Optimization results/` contains the results of our run of the optimization as configured above, plus a secondary run that incorporated more configuration tests for the LAP tracker. These results can be browsed with the command *Plugins > Tracking > TrackMate Helper results inspector.*

The optimum found by the procedure corresponds to a high score for each individual metrics. The downstream analysis focuses on quantifying morphological defects, and alteration of the cell cycle lengths, so we prioritize segmentation accuracy (SEG) and cell cycle accuracy (CCA) in our algorithm selection. SEG and CCA are both metrics whose values range from 0 to 1, 1 indicating a perfect score. The optimization identifies the *Omnipose detector* and the *Overlap tracker* as optimal on the tracking ground-truth we produced (Table 1). However, by inspecting the optimization results files, we find that there are many tracking configurations with good accuracy. For instance, over the 2590 valid configurations, 146 are found with a CCA value which is above 95% of the optimum. They include tracking configurations based on the *Omnipose* detector, and on the *Overlap* and *Trackastra* trackers.

These metrics are benchmarked against a ground truth derived from Omnipose segmentation outputs with manual refinement. This introduces an expected bias toward Omnipose, which would be problematic for algorithm ranking. However, since we visually confirmed that Omnipose could readily capture the bacteria shape and are not comparing algorithms but simply ensuring robust segmentation, this bias is acceptable. Notably, tracking based metrics such as CCA remain unbiased, as we reviewed and corrected all tracking errors. The tracking parameters that optimize both SEG and CCA are different. For downstream analysis, we select the one that maximizes the product of the SEG and CCA optimal values, which is the CCA optimum (0.882 vs 0.897).

| TrackMate helper optimization | Optimum for SEG | Optimum for CCA |
|---|---|---|
| Detector | **OMNIPOSE_DETECTOR**<br>- omnipose model: *bact_phase_omni*<br>- simplify contours: *true*<br>- cell diameter: *3.0* | **OMNIPOSE_DETECTOR**<br>- omnipose model: *bact_phase_omni*<br>- simplify contours: *true*<br>- cell diameter: *3.0* |
| Tracker | **OVERLAP_TRACKER**<br>- iou calculation: *PRECISE*<br>- scale factor: *1.2*<br>- min iou: *0.1* | **OVERLAP_TRACKER**<br>- iou calculation: *PRECISE*<br>- scale factor: *1.6*<br>- min iou: *0.3* |
| Spot filter | **AREA**<br>> 0 $\mu m^2$ | **AREA**<br>> 0 $\mu m^2$ |
| Score | SEG: 0.904<br>CCA: 0.976 | SEG: 0.904<br>CCA: 0.992 |

Table 1: Tracking parameters that optimize the SEG and CCA metrics.

#### 3b. Cross validation of the optimal tracking configuration and correcting the optimum

The optimum was obtained by running the optimization procedure on a movie following the WT strain. This movie is not in the dataset that will be used in the downstream analysis and was acquired with a different framerate (3 min vs 2 min). Additionally, we ran the optimization on a single movie that followed the WT strain. The mutant strains may introduce changes in morphology and dynamics that might cause tracking results obtained with this optimum to be incorrect. We need to validate that we can get good tracking results at least on a movie imaging a mutant. In this section we describe how we performed this validation. The corresponding files can be found in the `/3ter-CrossValidation` folder.

##### On UbiH mutant

All files are in the folder `/3ter-CrossValidation/UbiH` folder.

We first created a movie excerpt from the UbiH_M9glyCAAUra mutant movie by cropping a portion of the source image (`Excerpt_UbiH_M9glyCAAUra.tif`). We then generated a tracking ground truth for it, using the same approach as above (`Excerpt_UbiH_M9glyCAAUra-trackingGT_DONE.xml`). From this TrackMate file, we exported the ground-truth to the CTC file format (the `01_ST` folders).

We then ran TrackMate on the movie excerpt using the optimum configuration and saved the results (`Excerpt_UbiH_M9glyCAAUra-automaticTrackingResults.xml`). The plugin *Plugins* > *Tracking* > *TrackMate metrics computation* can be used to compute metrics on a single result. In this plugin panel we selected the CTC tracking metrics and entered the path to the automatic tracking results file and to the ground truth folder.

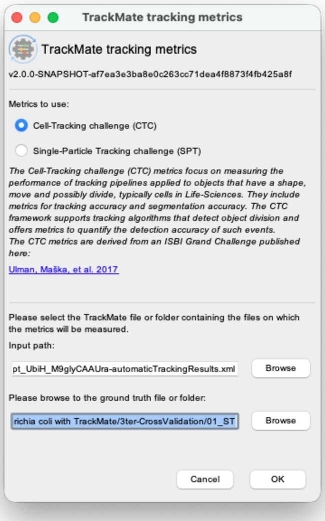

The plugin returns the following values:

```
- SEG          : 0.984
- TRA          : 0.982
- DET          : 0.982
- CT           : 0.805
- TF           : 0.989
- CCA          : 0.871
- BC           : 0.873
```

Clearly, the CCA metrics are significantly lower than what was obtained with the optimization movie. A cell-cycle accuracy value of 87% might be sufficient for downstream analysis, but it prompts an investigation. After inspecting the automated tracking results, we found that they contain many spurious tracks generated by small particles that were incorrectly detected by Omnipose and not filtered out. The parameter sweep in the first optimization included a sweep over a filter on spot area, but as the first movie did not contain these structures, the optimization did not select a filter value larger than 0 µm$^2$.

We ran a secondary optimization procedure on the UbiH_M9glyCAAUra excerpt movie, narrowing the parameter ranges to sensible value around the first optimum (`/3ter-CrossValidation/SecondaryOptimization`). Importantly, we kept the parameter sweep on the area filter. We found the following optima:

| TrackMate helper optimization | Optimum for SEG | Optimum for CCA |
|---|---|---|
| Detector | **OMNIPOSE_DETECTOR**<br>- omnipose model: *bact_phase_omni*<br>- simplify contours: *true*<br>- cell diameter: *3.0* | **OMNIPOSE_DETECTOR**<br>- omnipose model: *bact_phase_omni*<br>- simplify contours: *true*<br>- cell diameter: *3.0* |
| Tracker | **OVERLAP_TRACKER**<br>- iou calculation: *PRECISE*<br>- scale factor: *2*<br>- min iou: *0* | **OVERLAP_TRACKER**<br>- iou calculation: *PRECISE*<br>- scale factor: *1.6*<br>- min iou: *0.3* |
| Spot filter | **AREA**<br>> 0 µm$^2$ | **AREA**<br>> 1 µm$^2$ |
| Score | SEG: 0.989<br>CCA: 0.622 | SEG: 0.922<br>CCA: 0.972 |

Table 2: Secondary optimization results on Excerpt_UbiH_M9glyCAAUra.tif.

The optimum for CCA is the same that for the `20230331_washed_XY1.ome-1_stabilized_cropped.tif` movie, but with a filter on spot area larger than 1 μm$^2$. The SEG and CCA values it yields are satisfactory, and we use it for the next steps.

#### On UbiG mutant

All files are in the folder `/3ter-CrossValidation/UbiG` folder.

We repeated the same validation procedure on the UbiG mutant for ground-truth generation and metrics test. We found a significantly lower CCA value with the new optimum:

```
- SEG        : 0.957
- TRA        : 0.963
- DET        : 0.965
- CT         : 0.819
- TF         : 0.966
- CCA        : 0.612
- BC         : 0.880
```

Upon inspection, we identified 18 oversegmentation errors that incorrectly generated false cell division events. Although these errors represent a small fraction of the total objects in the movie (~8,500) they have a strong impact on the CCA value in the UbiG case. Indeed, the cell cycle length can only be measured between two cell divisions. For the UbiG mutant, we note that cell divisions are scarce. The movie `Excerpt_UbiG_M9glyCAAUra.tif` follows 83 lineages, but only 15 have more than 2 cell divisions, for a total of 99 division events in the whole movie. Most bacteria either divide once or fail to divide at all before growth arrest. By contrast, the wild-type (WT) movie analyzed earlier contains roughly 700 divisions across 72 lineages. The UbiG mutant actually displays cell cycle arrest, and the cell cycle length is therefore not meaningful for this mutant. We will incorporate this fact in the downstream analysis and use for the next steps the optimum for CCA found with the UbiH cross-validation.

### 4. Batch tracking

Step #4 in Figure 2.

Batch tracking requires a TrackMate file that contains the settings you want to use for batch processing. You can create it by running TrackMate on any movie and saving the results, or directly in the TrackMate helper. To do so, in select the *Best params* tab, and in it, select the *Best detector and tracker tab*. In the table there, select on the Segmentation accuracy line and click the *Launch TrackMate with selection* button. This opens the TrackMate plugin, preconfigured with the tracking parameters of the selected optimum. Click on the *Save* button at the bottom left of the TrackMate window to create a TrackMate file with these parameters and save it somewhere. You will find the settings resulting from the cross validation above in the file `/3ter-CrossValidation/ UbiH/ Excerpt_UbiH_M9glyCAAUra-automaticTrackingResultsWithAreaFiltering.xml`.

Close the image, the TrackMate-Helper window, the TrackMate window, and run the *Plugins > Tracking > TrackMate Batcher* command. It is made of 4 panels. In the *TrackMate settings* panel (top-right), drag and drop the TrackMate XML file that you saved at the previous step. The parameters of this tracking configuration should be displayed. In the `/4-Execute tracking in batch/` you will find the three image files of the *E. coli* dataset. Drag and drop them in the *Input images* (top-left) panel. Finally, in the *Outputs* panel (bottom-left), click the *TrackMate file (XML)* checkbox. Click the *Run* button (bottom-right) to run batch tracking. It should take about 5 - 10 minutes per image.

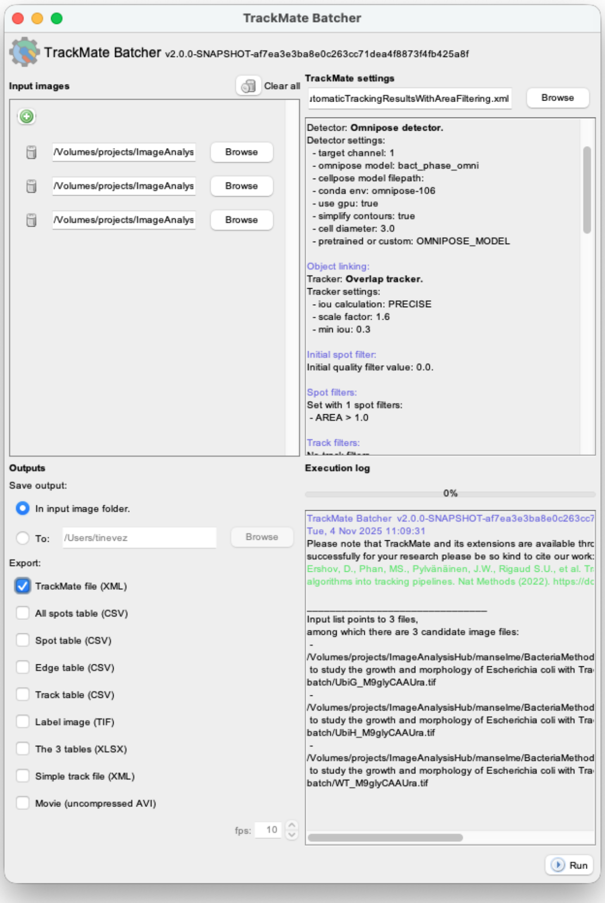

#### 5. Track analysis

Step #5 in Figure 2.

With the previous step, the tracking part of the protocol is over. It yields a set of tracking results that are optimal in the sense given above. Further analysis stems from this step, with code that is specific to each bacterial dynamics studies. We exemplify such an analysis code with the quantification of bacterial growth and the detection of morphology defects in mutants. You will find in the `/5-Analyze tracking results/` folder a Jupyter notebook that performs this analysis with pycellin [4]. The code copies the content from the github repository that contains python analysis code for all the examples of this article. It also contains installation instructions that we refer you to:

https://github.com/Image-Analysis-Hub/BacteriaMethodsCode

Open the file named `Ecoli_growth_and_morphology.ipynb` in Jupyter or VSCode, select the right Python environment, and run the notebook. It should complete in a few minutes and generate the raw figures below. This analysis is commented on in the main text, and we list below each of its steps.

Our protocol first identifies growth defects for UbiH and UbiG compared to WT. Cell divisions start only around frame 20 but are delayed and slowed down for UbiH. In the case of UbiG, it seems to halt after one or two divisions. For WT, the growth initially resembles an exponential growth, then becomes linear by the end of the movie, probably reflecting nutrient exhaustion.

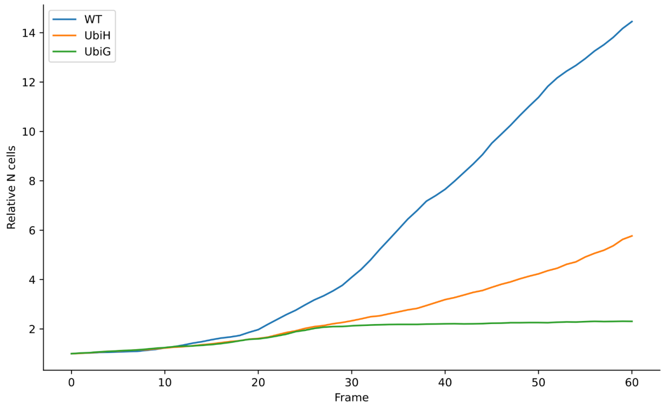

We then measure the cell cycle lengths. To avoid incorporating data that underestimate the cell cycle length or contain tracking mistakes, we filter out track segments that do not have exactly one parent (the division of the mother cell happened before the movie start or tracking mistakes) and exactly two daughters (the division happened after the movie end or tracking mistakes). Note that the displays of Figure 1 and Supplemental Movie 1 show unfiltered data. The distribution of cell cycle lengths highlights several subpopulations in the mutants. WT bacteria divide in about 35 minutes. The UbiH mutant has a large cell population that divides much slower. For the UbiG mutant, cell divisions are much more spread (Table 2), with populations with short cell cycle length and long ones. The results show marked and significant differences between WT and UbiH. Since we used a tracking configuration that gives an error on the cell cycle length of about 3% (CCA > 0.97), the difference we observe is likely not to be caused by tracking mistakes.

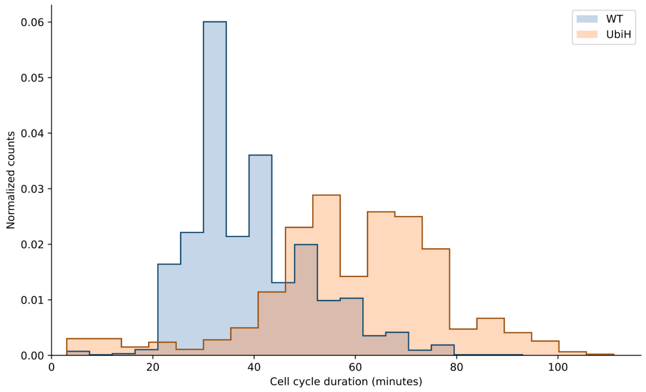

```
DESCRIPTIVE STATISTICS
=====================
        WT: 38.57 ± 11.81 min (N = 2139)
      UbiH: 59.84 ± 17.86 min (N = 860)

ONE-WAY ANOVA RESULTS
=====================
F-statistic: 1454.0428
P-value: 0.0000

TUKEY'S HSD POST-HOC TEST RESULTS
=================================
 Multiple Comparison of Means - Tukey HSD, FWER=0.05
=====================================================
group1 group2 meandiff p-adj  lower    upper   reject
-----------------------------------------------------
  UbiH     WT -21.2778   0.0 -22.3719 -20.1837   True
-----------------------------------------------------
```

Table 2: Comparison of cell cycle length duration.

The UbiG mutant quickly halts its growth. Most colonies have only one cell division. The numbers of cell divisions in WT and the UbiH mutant reflect the observed cell cycle length.

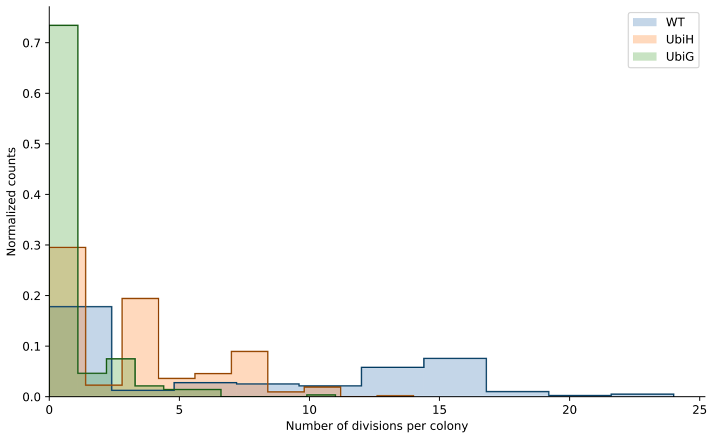

The morphology of the bacteria is also affected but differently depending on the mutant. The WT strain exhibits sometimes bacteria that are long (more than 6 µm), typically before cell division. The number of long bacteria stays small and constant throughout the movie. The UbiH mutant however produces an increasing number of abnormally long cells. In the case of UbiG the halting of bacterial growth is not accompanied by the appearance of morphological defects.

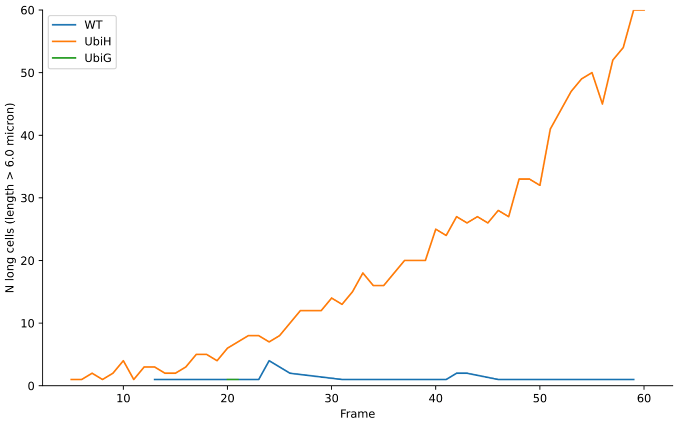

### B. Protocol to study the impact of mutations on the motility of *Helicobacter pylori*

*Helicobacter pylori* is a flagellated bacteria that can cause gastric ulcers. Its high motility, granted by the flagellum, is an important factor for its virulence [5]. In this protocol we will establish a pipeline to detect subtle changes in the motility of *H pylori* mutants, imaged in time lapse microscopy. This protocol will enable researchers to unravel the molecular mechanisms that drive *H pylori* movements.

#### Dataset and strategy

Download the dataset for this use-case can be downloaded here: DOI 10.5281/zenodo.17911259
https://doi.org/10.5281/zenodo.17911259

It is made of 9 long, high frame-rate movies with a rather large field of view that follow the motility of the bacteria. The 9 movies are split in 3 groups:

- Video 1-3: WT group. These bacteria are not transformed.
- Video 4-6: mutant group. The bacteria have two punctual mutations in genes thought to affect motility and virulence.
- Video 7.9: control group. These are the WT bacteria transformed with the antibiotic resistance cassette used for mutant selection.

In our experiments, we observed that there are fewer bacteria in the mutant group compared to the other two. CFU-based growth kinetics were conducted post-microscopy to elucidate density discrepancies between mutant and WT *H. pylori*. While OD curves overlapped, CFU counts showed a ~0.5-generation delay in the mutant (generation time = ~4 h), potentially explaining the observed lower number of bacteria in the movies of this group.

The goal of the following is to establish a protocol that can automatically measure motility features in these movies and perform comparative analysis between the groups. With such a protocol we should find that the motility features are similar between the control and the WT groups, and that the mutant group is different from these two. Additionally, we want to isolate what are the specific features that differentiate the mutant from the control groups, to learn how the motility is affected by the mutations.

The bacteria were imaged in brightfield. Due to bacteria's free movement, their contrast with the background can invert, making them appear white in some frames and black in others. High bacterial density, rapid movement, and frequent trajectory crossings, along with spurious dirt and aggregates, make detection and tracking challenging. Since we are only interested in the bacteria trajectories, we do not need a segmentation approach, and can rely on detection algorithms, which are often more robust. And we need this detection to be robust against the spurious objects in the field of view. We selected YOLO [6], a deep-learning detection and classification algorithm popular in computer vision. It requires training a model for our task, but we notice it displays good detection performance for a reasonable amount of annotation.

#### 0. Training a custom YOLO model for bacteria detection

In the protocol, this would be a step #0, before the steps described in Figure 2. This preliminary step is necessary because existing detection algorithms in TrackMate are unlikely to yield satisfactory results for our specific dataset.

##### Annotating images

Annotations were done on QuPath (version v0.5.1), but using any other software enabling the creation of bounding box annotations would also work. We randomly selected a set of frames extracted from the *H pylori* movies, ensuring that all the conditions (WT, mutant, control) were equally represented. We annotated these frames in QuPath by drawing rectangular bounding boxes around *H pylori* cells and assigning these annotations to a class named “cell”. A groovy script was then used to export these annotations at the YOLO format, and a dataset was created from a notebook to be compatible with the input expected by YOLO. More precisely, the dataset was built with 87 training images, 25 validation images and 14 test images (corresponding to 70% of the images for training, 20% for validation and 10% for testing). It can be found in the `/0-Train a custom YOLO model/` folder.

#### Training a YOLO model

We then initialized a YOLO v11 model with weights of the model *yolo11n*, pre-trained on the COCO dataset. We retrained it on our dataset for 300 epochs. The training configurations and data are available in an `hpylori_dataset.yaml` file and the `Train_yolov11_from_COCO_and_evaluate.ipynb` notebook in the `/0-Train a custom YOLO model/` folder. Evaluating this model on the test set enabled us to reach the following metrics:

| | |
|---|---|
| metrics / precision (B) | 0.83 |
| metrics / recall (B) | 0.85 |
| metrics / mAP50 (B) | 0.86 |
| metrics / mAP50-95 (B) | 0.62 |
| fitness | 0.64 |

The values of these metrics are not close to 1, indicating that the complexity of the background challenges the model's accuracy. Nonetheless, this will be enough for our tracking pipeline and get bacteria tracks long enough for the downstream analysis. It is also important to remember here that the precision and recall metrics are computed for fixed IoU and confidence thresholds. Therefore, these two metrics strongly depend on the confidence threshold and IoU threshold that were fixed for the model evaluation. More precisely, increasing the confidence threshold will reduce the false positive rate, increasing the precision. On the contrary, after increasing the confidence threshold, more true positives may be missed, thus reducing the recall. On the other hand, increasing the IoU threshold will require tighter overlaps between predicted and ground truth bounding boxes to be considered as positive detections, leading to fewer predictions classified as true positive and thus decreasing the recall. The metrics were computed for a confidence threshold of 0.1 and IoU threshold of 0.4. By definition, the three other metrics computed here are either computed over a range of IoU threshold values, or on an explicit IoU threshold value, making them more relevant to compare different models.

### 1. Generate an initial result for a tracking ground-truth

Step #1 in Figure 2.

We will use the same strategy as in the previous use case. After choosing one movie we will use TrackMate with the YOLO model we just trained, and a sensible choice for a tracker. We will then fix tracking mistakes manually in TrackMate.

First install the required TrackMate extension, as explained here: https://imagej.net/plugins/trackmate/detectors/trackmate-yolo

To accelerate annotations, we will generate a ground-truth for a subset of the first movie of the dataset.

- Open Fiji. In the folder `/1-Generating an initial results for a tracking ground-truth` you will find a cropped version of the first movie, over 60 frames and 1024x1024 pixels named `SHicham_Video1_crop.tif`. Open it.
- Launch TrackMate from the menu *Plugins* > *Tracking* > *TrackMate.*
- Click the *Next* button in the TrackMate user interface (UI). The panel '*Select a detector*' is shown. In the list, select '*YOLO detector*' then click *Next.* If this item is not present in the list, you need to install the corresponding extension, as explained in the installation section above.
- The '*YOLO detector*' panel is shown. There are only a few parameters:

- The ‘*Conda environment*’ is important. It should be set to the conda environment in which you installed YOLO. If it is not listed, follow the instructions in the TrackMate-YOLO documentation page listed above.
- For the ‘*Path to a YOLO model*’, browse to the results of the training step documented above. If you skipped it you can find the model used to generate the results in the main text in the folder `/0-Train a custom YOLO model/results/best 8.pt`.
- You can leave ‘*Confidence threshold*’ and ‘*IoU threshold*’ to their default values (0.25 and 0.7).

Click the *Preview* button. The detection should run without error, and you should get decent detection results. If not, the installation of YOLO and its TrackMate integration probably did not work. Follow the instructions at the beginning of this section. Note that the detection results are represented as TrackMate circular spots. This YOLO integration does not yield the object shape (which we do not need) and there is no object contours returned.

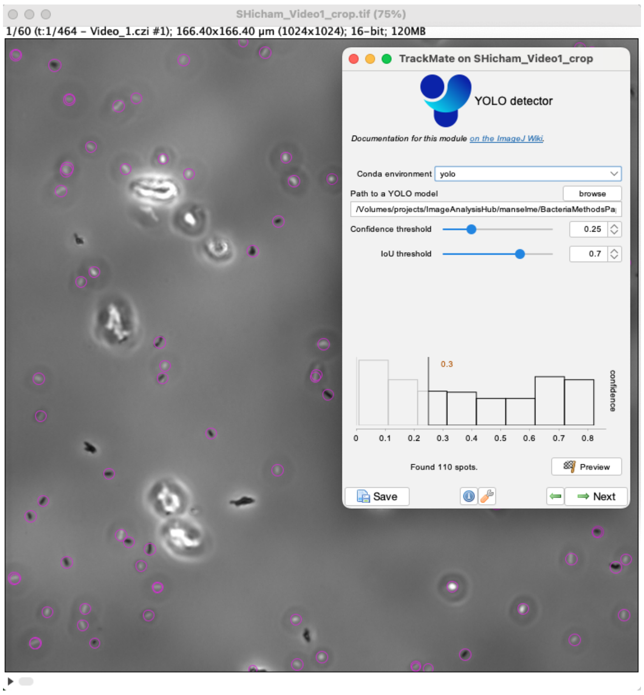

- Click *Next*. The detection runs over all the 60 frames, in about 1 minute depending on your workstation.
- Click *Next* again. Make sure that the *Initial thresholding* step does not filter out any spots, and there are no spot filters in the next panel.
- Click *Next* again and in the tracker selection panel, pick the Kalman tracker. This is a reasonable choice in our case, as the bacteria do not diffuse but move in a nearly constant direction.
- When configuring the Kalman tracker, use 5 µm for the ‘*Initial search radius*’, 5 µm for the ‘*Search radius*’ and 0 for the ‘*Max frame gap*’. We will anyway fix the gaps manually by adding the missed detections later.
- With these parameters you will get a starting point for the manual corrections of the tracking results. Save it.

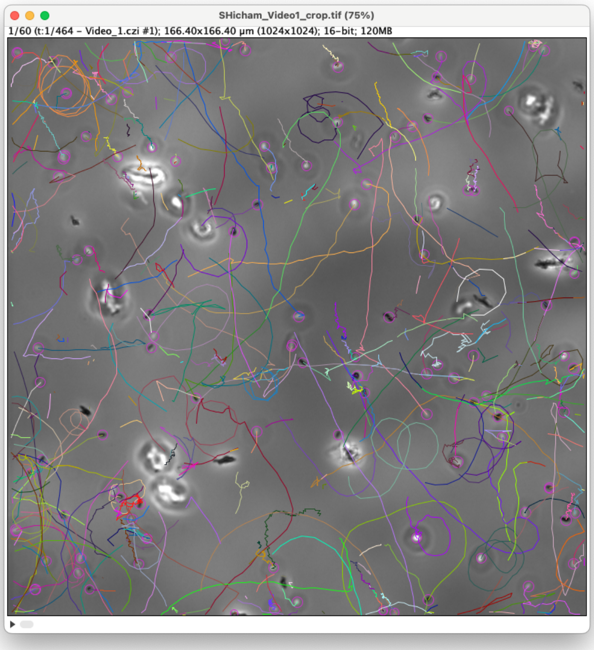

#### 2. Curating the automated tracking results

Step #2 in Figure 2.

This time the manual correction will not involve editing the segmentation results. We just need to have exactly one detection, in the shape of a circular spot, per bacteria. We also do not aim at tracking bacteria when they go through dirt or aggregates and should remove any detection on these spurious objects. Finally, we want to ensure the tracks we get effectively come from single bacteria and should delete tracks for moving objects that are sure to be made of more than one bacterium.

The full correction can take several afternoons. The result from our corrections, used in the main text, can be found in the `/2-Curating the automated tracking results` folder under the name `SHicham_Video1_crop-GT_DONE.xml`

#### 3. Optimizing tracking parameters

##### Configuration

Step #3 in Figure 2.

We will be using the TrackMate-Helper again, but this time using different metrics than the CTC metrics used in the previous use case. Indeed, the bacteria we track here are not dividing, and we are using only their trajectories for the downstream analysis. The adequate metrics for this endeavor are the Single-Particle Tracking (SPT) challenge metrics [2].

Launch Fiji and open the image named `SHicham_Video1_crop.tif` in the folder `/3-Optimize tracking parameters with the TrackMate-Helper`. Launch the TrackMate Helper in *Plugins > Tracking > TrackMate Helper*. In the panel that appears, select *Single-Particle Tracking challenge (SPT)* for the metrics. A text box appears, prompting for a *Max distance for pairing*. In the SPT metrics, detection results are 2D and 3D points. To assess whether they match a detection in the ground-truth, the metrics calculation simply takes the closest one. If it cannot find one within the specified *Max distance for pairing*, it is considered a false positive. If a point in the ground truth does not have a detection result within this distance, it is considered a false negative. You need to specify this max distance here in the units used in the source image. In our case it is µm, and we can use a max pairing distance of 1 µm, which is reasonable given the size of the bacteria we track. In the image list, select the `SHicham_Video1_crop.tif`. And in the *ground truth file or folder path*, browse to the `SHicham_Video1_crop-GT_DONE.xml` which is a copy of the tracking ground-truth TrackMate file generated in the previous step. Click *OK* and the helper configuration panel should open.

In the folder of `SHicham_Video1_crop-GT_DONE.xml` we prepared a configuration with a sensible choice of algorithms, filters and parameters to choose, and that will be picked up when you open the helper. You still need to edit the conda environment for the YOLO and Trackastra modules and point the *Path to a YOLO model* to the model you trained, or the one we prepared in the folder `/0-Train a custom YOLO model/results/best 8.pt`. You can see that we aim at testing across more than 100k different configurations. Fortunately, the SPT evaluation is very fast compared to the CTC metrics, and this can be done overnight.

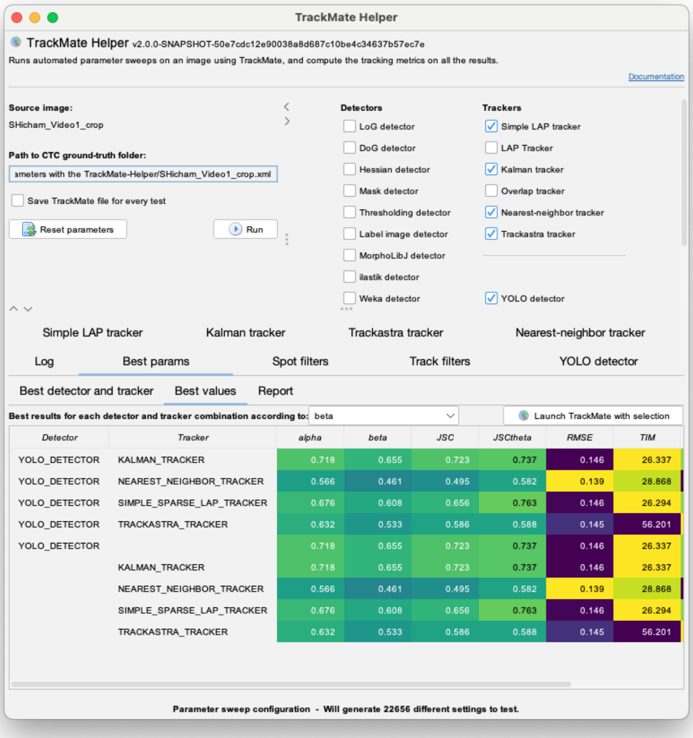

Given the complexity of the image background, we are likely to get false positive detections, and they might create short spurious tracks. We can optimize against those by adding a parameter sweep on track filters, filtering out tracks that are static. This is what is done in the *Track filters* tab of the helper.

#### Interpreting optimization results

As the downstream analysis involves computing track features (straightness, mean speed, etc.) to investigate how bacteria motility changes with mutations, the tracking pipeline must be optimized with respect to matching the ground truth and penalizing spurious tracks. The best parameter for this is the SPT '$\beta$' metric that computes the matching score between tracks and the ground truth and penalizes false positive tracks. Its definition is detailed in [2] ("Supplementary Note 3: Performance Measures") that we summarise here. First, the $\alpha$ metric is defined as follow:

$$\alpha = 1 - \frac{d(X,Y)}{d(X,\emptyset)}$$

where $d(X,Y)$ is the distance from the set of ground truth tracks ($X$) to the estimated tracks ($Y$). This distance is defined by pairing the sets of estimated versus ground truth tracks, then summing the distance of individual tracks. For one track, the distance is defined as the sum of Euclidean distances between each detected object and its corresponding ground-truth match, clipped at a maximum threshold $\varepsilon$ (the max pairing distance in the helper above). The normalization term $d(X,\emptyset)$ is the distance value if all objects of all estimated tracks were farther than $\varepsilon$ from any object in the ground truth tracks. It is an upper bound for $d(X,Y)$. With this definition, $\alpha$ ranges from 0 to 1. A value of 1 indicates perfect alignment between estimated and ground-truth tracks, both in spatial localization (object positions) and temporal extent (track duration). Conversely, $\alpha$ = 0 signifies a complete mismatch. Importantly, we see that the $\alpha$ metric compounds tracking accuracy (are the tracks complete, or stop too early, or are they missing objects, or do they conflate several ground truth tracks?) with object localization accuracy (how far is an object from the ground truth position?) The SPT metrics were originally developed to analyze trajectories of sub-resolved particles. In this work, we adapt these metrics to complex objects such as bacteria, where the true center position is ill-defined and highly sensitive to detection noise and segmentation errors. We therefore expect the value of $\alpha$ to be low because of localization accuracy even if estimated tracks match the ground truth within the tolerance defined by $\varepsilon$.

The definition of $\beta$ is built on $\alpha$ but adds a penalization for false positive tracks:

$$\beta = \frac{d(X,\emptyset) - d(X,Y)}{d(X,\emptyset) - d(\bar{Y},\emptyset)}$$

where $\bar{Y}$ is the set of spurious tracks, and $d(\bar{Y},\emptyset)$ the penalty distance obtained by summing the distance gate $\varepsilon$ for all spurious detections. The values of $\beta$ range from 0 to $\alpha$ and reach the latter if there are no spurious tracks.

You will find the results of our run in the folder `/3bis-Optimization results`, where we have added the LoG detector to illustrate the metric degradation with an inadequate detector. To display and summarize them, you can use the Helper results inspector plugin in *Plugins* > *Tracking* > *TrackMate Helper results inspector.* Point to path to the results to this folder and select *SPT* as metrics. The following panel should appear.

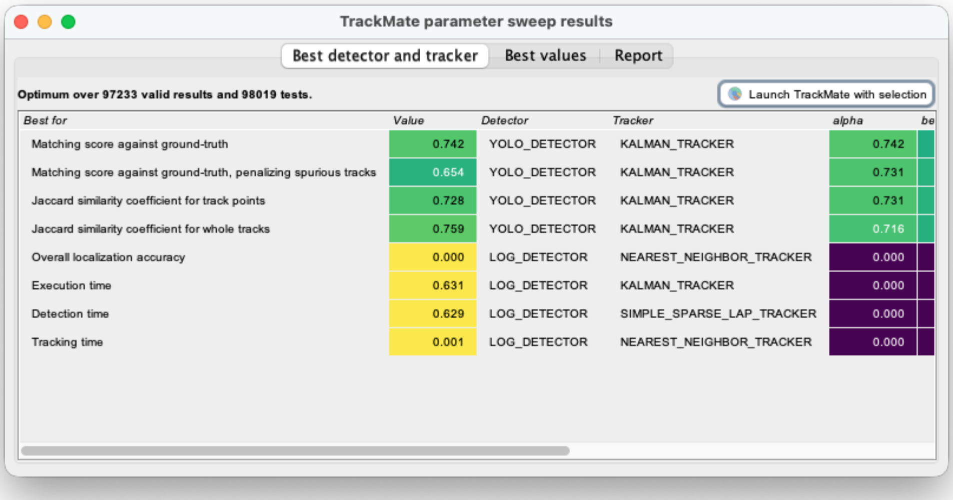

| Best for | Value | Detector | Tracker | alpha | be |
|---|---|---|---|---|---|
| Matching score against ground-truth | 0.742 | YOLO_DETECTOR | KALMAN_TRACKER | 0.742 | |
| Matching score against ground-truth, penalizing spurious tracks | 0.654 | YOLO_DETECTOR | KALMAN_TRACKER | 0.731 | |
| Jaccard similarity coefficient for track points | 0.728 | YOLO_DETECTOR | KALMAN_TRACKER | 0.731 | |
| Jaccard similarity coefficient for whole tracks | 0.759 | YOLO_DETECTOR | KALMAN_TRACKER | 0.716 | |
| Overall localization accuracy | 0.000 | LOG_DETECTOR | NEAREST_NEIGHBOR_TRACKER | 0.000 | |
| Execution time | 0.631 | LOG_DETECTOR | KALMAN_TRACKER | 0.000 | |
| Detection time | 0.629 | LOG_DETECTOR | SIMPLE_SPARSE_LAP_TRACKER | 0.000 | |
| Tracking time | 0.001 | LOG_DETECTOR | NEAREST_NEIGHBOR_TRACKER | 0.000 | |

And in the *Report* tab, you will find the parameters that maximize $\beta$ are:

```
________________________________________________________________________
Best configuration for Matching score against ground-truth, penalizing spurious tracks with a
score of 0.654
For detector: YOLO_DETECTOR with settings:
  - conda env: yolo
  - yolo model filepath: ./0bis-Train a custom YOLO model/results/best 8.pt
  - yolo iou threshold: 0.30000000000000004
  - use gpu: true
  - yolo conf threshold: 0.1
And tracker: KALMAN_TRACKER with settings:
  - max frame gap: 4
  - kalman search radius: 5.0
  - linking max distance: 3.0
```

```
With track filters:
 - MAX_DISTANCE_TRAVELED > 5.0
Single-Particle Tracking (SPT) Challenge metrics metrics:
 - alpha         : 0.731
 - beta          : 0.654
 - JSC           : 0.728
 - JSCtheta      : 0.725
 - RMSE          : 0.151
 - TIM           : 22.949
 - DETECTION_TIME: 22.676
 - TRACKING_TIME : 0.273
```

We note that the score we get for $\beta$ comparatively low (65%). As we noted above, this metric is very sensitive to deviation from the ground truth, and values close to 50% are common when the density of objects is medium to high (Figure 2 in [2]). By inspecting the results, we see that some tracks have gaps when a detection is missed. These tracks still extend over time, as the Kalman tracker can bridge over gaps, but the missed detections are still penalized in the β score. Conversely the YOLO detector sometimes creates multiple detections for a single bacterium, which results in several tracks per bacterium, lowering the β score as well. We therefore need to validate whether this optimum can yield meaningful and robust measurements based on the tracks it estimates.

### 3b. Cross validation and validation

We first cross-validate the optimum on the most different group in the dataset. In our case, this is the mutant group. As for the previous use case, we generated a tracking ground truth for an excerpt of a movie of this group. You can find the movie and TrackMate file in the folder `/3ter-CrossValidation/mutant`. We then ran automated tracking with the parameters of the optimum and measured the tracking metrics of the results with the *Plugins* > *Tracking* > *TrackMate metrics computation* plugin. We get the following results:

```
- alpha         : 0.655
- beta          : 0.627
- JSC           : 0.642
- JSCtheta      : 0.600
- RMSE          : 0.122
```

The value for $\alpha$ and $\beta$ are commensurate with that of the WT movie.

These relatively low scores are not inherently problematic. Our downstream analysis relies on track-averaged features like confinement ratio and mean speed, which are robust to localization errors and pointwise tracking mistakes. To verify this robustness, we validate our approach by comparing analytical results derived from ground-truth tracks against those obtained using our optimized tracking method. The data and script that performs the validation can be found in `/3ter-CrossValidation/validation/`. Briefly, we found out that there are no significant differences between the track features measured on the ground truth tracks versus the estimated tracks, except for the *total distance travelled* feature.

### 4. Batch tracking

Step #4 in Figure 2.

Run the full tracking pipeline with the optimal parameters on the movie excerpt and save the TrackMate XML file resulting from this. We now need to perform tracking with these optimal parameters over the 9 movies of the dataset. For this we will use the TrackMate bather in *Plugins* > *Tracking* > *TrackMate Batcher*, as described in the previous section.

Once the batcher window is opened, drag and drop the 9 movies in `/4-Batch tracking with TrackMate` in the *Input images* top left quadrant. Additionally, add the Drag and drop the TrackMate file with optimal parameters you saved in the previous paragraph in the top right *TrackMate settings* quadrant (you can find it in `/4-Batch tracking with TrackMate/optimum-tracking-pipeline.xml`). In the *Output* quadrant, select a folder where to save results, and check *TrackMate file (XML)* and *Track table (CSV)*. Then click the *Run* button. The 9 movies are larger and longer than the one used to create the ground truth, and the total processing will take 30 min - 1 hour. In the `/results` subfolder you will find the results of our run. It contains two files per movie, the TrackMate XML and the track CSV files, as we specified.

### 5. Track classification

Step #5 in Figure 2.

With the previous step, the tracking part of the pipeline is finished. We need now to analyze tracks and make sense of the motility we measured. Here we chose to directly rely on the numerical track features measured in TrackMate and stored in the CSV files we created. You will find that each of these CSV files is a table containing one row per track, with various track measurements made in TrackMate (‘`LINEARITY_OF_FORWARD_PROGRESSION`’, ‘`TRACK_DISPLACEMENT`’, etc.) We will interpret the motility of *H pylori* in the movies based on these values only. As for the previous use case, we prepared the analysis scripts and the exported track data in the `/5-Analyze tracking results/` folder. It is a copy of the script maintained here:

https://github.com/Image-Analysis-Hub/BacteriaMethodsCode

Open the file `Classify_h_pylori_TrackMate_tracks_by_group.ipynb` in a jupyter notebook or in VSCode for instance. Its header contains instructions to install dependencies with conda. The notebook also has annotations that explain the steps we took.

Briefly, we start by importing the CSV files in pandas data structures and clean the resulting tables from irrelevant or unsuited features (like track name, id, track position, number of spots in the track). We then measure cross-correlation of the remaining features and use it to remove redundant features.

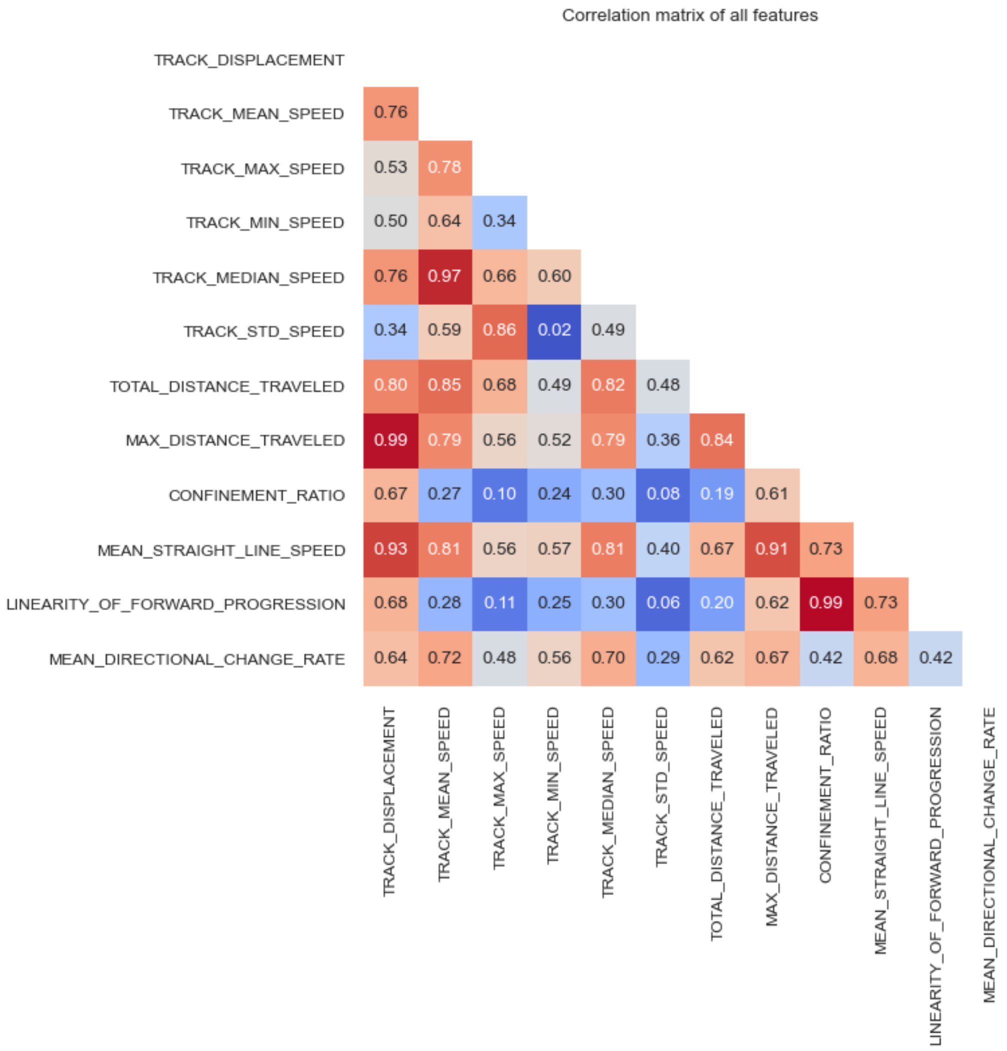

Without surprise, we find that features such as MAX_DISTANCE_TRAVELED correlate with TRACK_DISPLACEMENT and LINEARITY_OF_FORWARD_PROGRESSION correlate with CONFINMENT_RATIO. By setting a threshold of 80% we end up in pruning the following features:

- LINEARITY_OF_FORWARD_PROGRESSION
- MAX_DISTANCE_TRAVELED
- MEAN_STRAIGHT_LINE_SPEED
- TOTAL_DISTANCE_TRAVELED
- TRACK_MEDIAN_SPEED
- TRACK_STD_SPEED

In a second step we compared the WT group and the control group. The control group has been transfected with a mock cassette, and this comparison will allow concluding on the impact of the transfection itself on bacterial motility. We find that the motility features of these two groups are not significantly different, suggesting that the transfection does not affect the motility. We then performed the same comparison between the control group and the mutant group and found that motility features are statistically different in these two groups. By testing for effect size (with Cohen's d value) we find that all differences are small or negligible. The main feature differences are for the bacteria speed.

### Supplemental Methods

#### Computers used

The training of the YOLO model was done on an Ubuntu workstation equipped with two NVIDIA RTX4090 GPU cards, 128 GB RAM and an AMD Ryzen 7950X CPU. The use cases detailed in the supplemental information were processed with a 2023 Mac Studio with an Apple M2 CPU and 64 GB RAM.

#### Software version

Several new versions of Fiji and TrackMate plugins were developed for this work and used for the use cases above: *TrackMate* v8.1.6, *TrackMate-Helper* v2.0.0, *TrackMate-Cellpose* v1.0.1, *TrackMate-Trackastra* v2.0.2, *TrackMate-YOLO* v1.0.1, all running in *Fiji* v2.16.1 ('latest' branch) with Java 21. The Python algorithms used with TrackMate are *Omnipose* v1.0.6, *Trackastra* v0.5.2 and *YOLO* v8.3.85. For the step 5 of the use cases, we used the following libraries running with Python 3.10: *pycellin* v0.4.2, *scikit-image* v0.25.2, *scikit-learn* v1.7.2, *numpy* v2.2.6, *matplotlib* v3.10.8, *seaborn* v0.13.2.

#### Imaging *E. coli* growth

*E. coli* strains were cultured in M9 medium supplemented with 0.4% glycerol 0,1% CAA and 20ug/mL uracil at 37°C as described in [7]. For imaging cells were immobilized on 1% agarose pads supplemented with same medium and imaged using 100 ms phase acquisition in time-lapse series. Images were acquired with a fully motorized Nikon Ti-2E Eclipse inverted microscope at 37°C equipped with a CFI Plan Apochromat λ DM 100XH 1.45/0.13 mm Ph3 oil objective (Nikon), a SpectraX illuminator (Lumencor), an Orca-Flash 4.0 V3 camera (Hamamatsu). Multi-dimensional image acquisition was made using NIS-Ar software (Nikon). Microscopy data reported in this paper is included in the dataset linked in the first use case.

#### Imaging *H. pylori* motility

*H. pylori* strains were grown in Blood agar plates with 10% defibrinated horse blood and the following antibiotics-antifungal mixture: amphotericin B 2.5 μg/mL, polymyxin B 0.31 μg/mL, trimethoprim 6.25 μg/mL, and vancomycin 12.5 μg/mL. For liquid cultures, we used Brain Heart Infusion (BHI) broth (Oxoid) supplemented with 10% Fetal Calf Serum (Eurobio) and the antibiotics−antifungal mixture. *H. pylori* cells were grown at 37°C under microaerophilic conditions (6% O2, 13% CO2, 81% N2) using an Anoxomat (MART Microbiology) atmosphere generator and at 180 rpm shaking. For imaging, cells were observed in exponential growth phase in same liquid media. Cells were added to a custom-made well in an agar pad to avoid fluid movement. Images were acquired with a Zeiss AxioObserver Z1 inverted microscope in a temperature and atmosphere-controlled chamber (37°C and 6% $O_2$, 10% $CO_2$, 84% $N_2$ saturated with water) with the Zen software package, an Orca-Flash 4.0 camera (Hamamatsu), and a LD Plan NeoFluar 40X/0.6 mm Koo PH2 M27 objective (Zeiss) in phase contrast. Time-lapse movies were obtained with 50 ms frame interval for 30 seconds.

### Supplemental Tables

#### Supplemental Table 1

| **Time estimates for each step** | **Use case 1: Impact of the respiratory chain in the growth and morphology of *Escherichia coli*** | **Use case 2: Impact of mutations on the motility of *Helicobacter pylori*** |
|---|---|---|
| *Preparing.*<br>*Installing Fiji, collating and organizing the data.* | < 1 hour | < 1 hour |
| *Step #0.*<br>*Training a custom detector for the specific dataset.* | NA.<br>We use a builtin model. | Train a custom YOLO model based on annotations done in QuPath.<br>- One session of 1.5 hours. Active work.<br>- training time on a workstation: 3 hours. Passive. |
| *Step #1.*<br>*Generating an initial result for a tracking ground-truth* | < 1 hour | < 1 hour |
| *Step #2.*<br>*Curating the automated tracking results* | 3 - 6 sessions of 2 - 3 hours. Active work. | 4 - 8 sessions of 2 - 3 hours. Active work. |
| *Step #3.*<br>*Optimizing tracking parameters with the TrackMate-Helper* | Overnight.<br>Passive. | Overnight (for 100k configurations tested) or 2-4 hours (~10k configurations).<br>Passive. |
| *Step #3ter.*<br>*Cross validation and validation* | Cross validation requires generating a second tracking ground-truth.<br>If the second movie excerpt has the same size:<br>3 - 6 sessions of 2 - 3 hours. Active work. | Cross validation requires generating a second tracking ground-truth, but here it was made on a movie with fewer bacteria (mutant group):<br>2 sessions of 1 - 3 hours. Active work.<br><br>Validation is based on comparing tracking outputs<br>< 2 hours. Active work. |
| *Step #4.*<br>*Batch tracking with TrackMate* | < 1 hour<br>However here the example dataset only includes 3 movies for demonstration purposes. Provision about 10 minutes per image.<br>Passive. | < 1 hour<br>Provision about 5 - 10 minutes per image.<br>Passive. |
| *Step #5.*<br>*Analyze tracking results* | Elaborating the analysis script: 1 - 2 days. Active work.<br>Running the analysis and collecting results: < 1 hour. Passive. | Elaborating the analysis script: 1 - 2 days. Active work.<br>Running the analysis and collecting results: < 1 hour. Passive. |

**Supplemental Table 1:** Estimated time required for each step in the two use cases presented in this study. *Active work* refers to tasks requiring direct, sustained effort by a scientist. *Passive* work refers to computational steps performed unsupervised by a workstation.

### Supplemental Movies Legends

#### Supplemental Movie 1

Time-lapse movies following the growth of the WT (left) and UbiH (right) *E. coli* strains in the first use case. Bacteria are overlaid with segmentation results color-coded for the current cell cycle length. Cell cycle lengths longer than 90 minutes are colored in dark red. On this display, the cell cycle lengths of bacteria that divided before the movie started or after the movie ended are underestimated, as the cell cycle lengths are measured from the beginning, respectively to the end, of the movie for these bacteria.

#### Supplemental Movie 2

Time-lapse movies following the growth of the WT (left), UbiH (center) and UbiG (right) *E. coli* strains in the first use case. Bacteria are overlaid with segmentation results color-coded for their length. Cells that are longer than 6 µm are colored in dark red.

#### Supplemental Movie 3

Combined movie that follows the motility of the three *Helicobacter pylori* strains used in the second use case. The results of automatic tracking with optimal tracking configuration are overlaid. Left: wild-type; middle: mutant strain; right: control strain.
